# PXC2, a polarized receptor kinase, functions to repress ground tissue cell divisions and restrict stele size

**DOI:** 10.1101/2021.02.11.429611

**Authors:** Jason Goff, R. M. Imtiaz Karim Rony, Zengxiang Ge, Jakub Hajný, Cecilia Rodriguez-Furlan, Jiri Friml, Jaimie M. Van Norman

## Abstract

Coordination of cell division and differentiation is critical for tissue patterning during organ development. Directional signaling and cell polarity have important roles in coordination of these processes. For instance, the Arabidopsis receptor-like kinase INFLORESCENCE AND ROOT APICES KINASE (IRK) functions to restrict stele area and repress longitudinal anticlinal divisions (LADs) in the endodermis where it is polarly localized. IRK is closely related to PXY/TDR CORRELATED 2 (PXC2); here, we show that PXC2 exhibits similar polarized accumulation suggesting they have related functions. *pxc2* roots have increased stele area and *irk pxc2* double mutant roots show increases in stele area and endodermal LADs beyond either single mutant, indicating that polarly localized IRK and PXC2 function redundantly to repress endodermal LADs and stele area. The double mutant also exhibits agravitropic root growth and abnormal cotyledon vein patterning. Manipulation of PIN1 trafficking and (re)localization mechanisms indicate that PXC2 and/or IRK suggest a link to modulation of polar auxin transport. We conclude that IRK and PXC2 have partially redundant functions in the root, but may have independent functions that are tissue-specific. We propose that repression of endodermal LADs and stele area through a PXC2/IRK-mediated, directional signaling pathway is required for normal ground tissue cell divisions, tissue patterning, and growth in Arabidopsis roots.

## INTRODUCTION

In plant development, proper orientation of cell divisions relative to the body axis is required for tissue and organ patterning and overall organ shape (Shao and Dong, 2016; Facette et al., 2018; Rasmussen and Bellinger, 2018; Martinez et al., 2017). Defects in tissue patterning and organ shape can negatively impact organ function (Meyerowitz, 1997; Camilleri et al., 2002). The organization of root tissues and cell types is critical to its function in anchorage and uptake and can be partly attributed to strict regulation of the orientation of root cell divisions. In the root, cell divisions are typically oriented periclinally (parallel to the root’s surface) or anticlinally (perpendicular to the root’s surface) which generates more cells in the radial and/or longitudinal axes. The highly stereotypical organization of root cell types and tissues facilitates investigation into the mechanisms underlying coordination of oriented cell division and tissue patterning during organ development.

In the Arabidopsis root, the central stele, which contains the pericycle and the vascular tissues, is surrounded by the endodermis, cortex, and epidermis in the transverse axis (Figure 1) (Dolan et al., 1993). Cell division orientation in the root is strictly controlled such that cells are maintained in linear files that extend from the stem cell niche shootward in the root’s longitudinal axis and, in the transverse axis, the number of cells in the rings around the stele is consistent for each cell type. For example, in the root ground tissue, the cortex and endodermal cell types are derived from a single stem (initial) cell, the cortex/endodermal initial (CEI). There are typically 8 CEI that divide anticlinally to produce a daughter cell which undergoes a periclinal cell division to produce a pair of cell types: the endodermis towards the inside and the cortex towards the outside (Figure 1A) (Dolan et al., 1993; Scheres and Benfey, 1999). Longitudinal anticlinal divisions (LADs), which would add more cells within a ring, rarely occur in the endodermis and cortex, such that typically there are eight of each of these cell types surrounding the stele. Following cell proliferation in the longitudinal axis of the root meristem, cells undergo dramatic elongation which accounts for the majority of root lengthening (Hodge et al., 2009). In order for the root to effectively navigate a dynamic and heterogeneous soil environment, precise cellular organization together with coordinated cell elongation must be maintained. Differential cell elongation within the root allows for directional (tropic) root growth in response to various environmental stimuli (Correll and Kiss, 2005; Dyson et al., 2014; Dietrich et al., 2017). Disruption of the root’s ability to coordinate cell division and/or cell elongation often leads to defects in root form and function.

**Figure 1.**
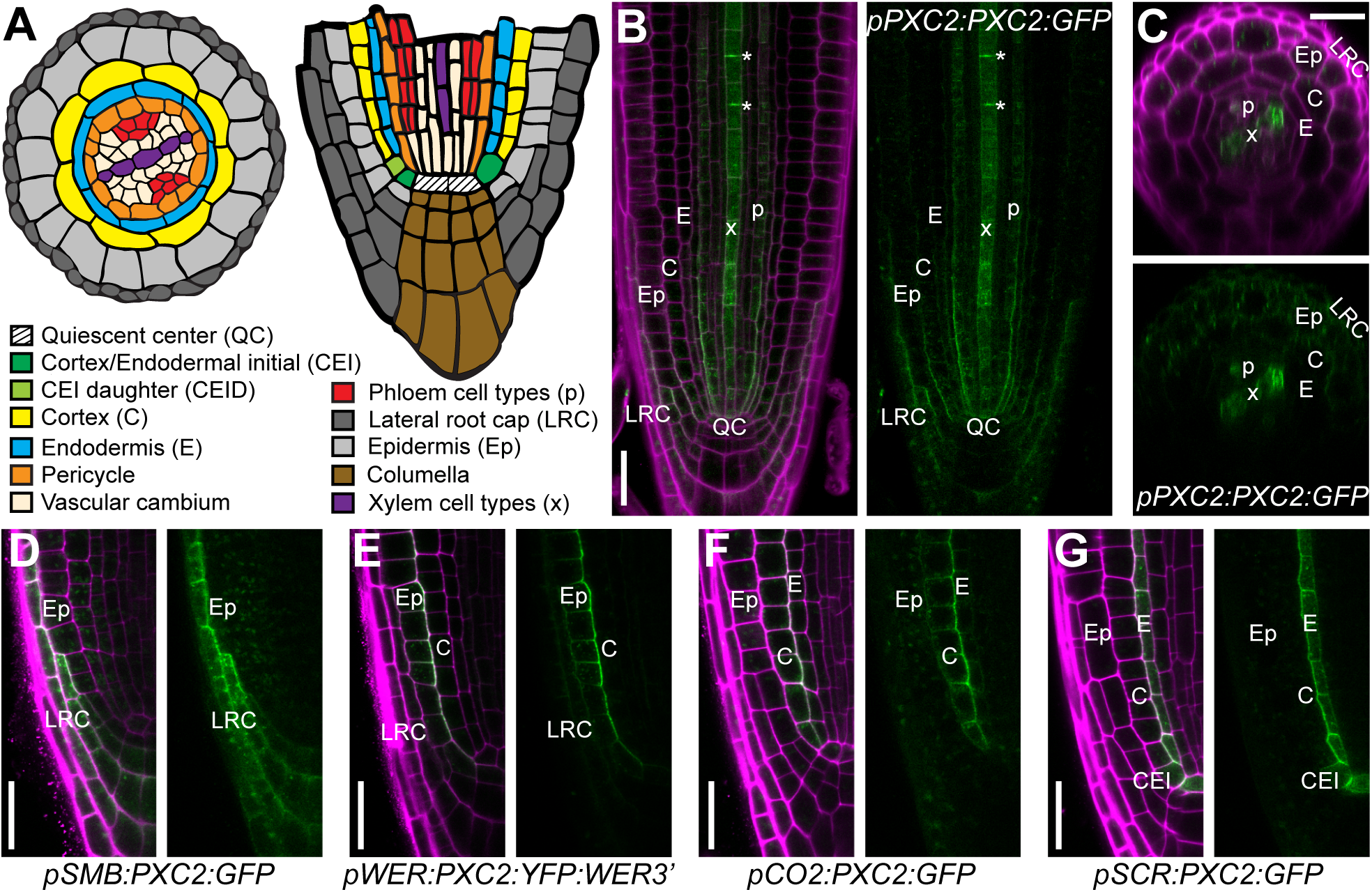
PXC2 is polarly localized in root cell types. (A) Schematics of an Arabidopsis thaliana root meristem with transverse (left) and longitudinal (right) sections showing cell types and their organization (adapted from (Campos, et al., 2020)). (B-G) Confocal micrographs of Arabidopsis root meristems stained with propidium iodide (PI, magenta) to visualize individual cells and showing FP (fluorescent protein) localization (green) with PI + FP (right) and FP alone (left). (B) Median longitudinal and (C) transverse optical sections showing PXC2-GFP accumula-tion when driven by *pPXC2*. (D-G) Polar localization of PXC2-FP upon misexpression by (D) *pSMB*, (E) *pWER*, (F) *pCO2*, and (G) *pSCR*. Cell type abbreviations as in panel (A) and asterisks (*) indicate recently formed cell plates during xylem cell division. Scale bars: 25 μm.

Coordination of cellular processes, including division and elongation, across cell types requires cell-cell communication (Van Norman et al., 2011; Wu et al., 2016; Chaiwanon et al., 2016). Furthermore, cell division orientation and cell fate specification cues are proposed to be largely informed by extrinsic factors. This model is supported by cell ablation studies in the root meristem that show altered cell division patterns and cell fate specification when cells are lost and replaced through division of internal neighbors (van den Burg et al., 1995; van den Berg et al., 1997; Marhava et al., 2019). In plants, a large family of receptor-like kinases (RLKs) is proposed to serve as major participants in intercellular communication across the plant body. Over 600 RLKs are encoded by the Arabidopsis genome and they are grouped into different subfamilies, such as leucine-rich repeat (LRR) RLKs, which contain multiple leucine-rich repeats in their ectodomain (Shiu and Bleecker 2001; Shiu and Bleecker 2003; Dievart et al. 2020). These LRR-RLKs are generally predicted to perceive extracellular ligands and activate downstream signaling pathways. However, despite decades of investigation, the function of many LRR-RLKs remains unknown partially due to high levels of functional redundancy and/or very specific or mild abnormal phenotypes (Diévart and Clark, 2003).

In the Arabidopsis root the LRR-RLK, IRK, is required to repress endodermal LADs and restrict stele area in the radial axis. These defects are rescued by expression of IRK from its endogenous promoter or by endodermal-specific expression of IRK, suggesting that abnormal LAD of endodermal cells is the primary defect in *irk* mutants (Campos et al., 2020). IRK is polarly localized in root cell types, with accumulation at the outer plasma membrane (PM) domain in the endodermis and pericycle and the inner domain in cells of the epidermis and cortex. Its polar distribution is informed locally by adjacent cells and achieved through polar secretion directed by sorting determinants in its extracellular domain (Campos et al., 2020; Rodriguez-Furlan et al., 2022). Despite the cell level abnormalities in *irk*, root growth and overall morphology appear largely normal. This prompted us to investigate whether the function of a related LRR-RLK was obscuring a more detrimental dysregulation of ground tissue patterning and an enhanced impact on root growth and development.

IRK is most closely related to PXY/TDR-CORRELATED 2 (PXC2, encoded by At5g01890) (Shiu and Bleecker, 2001, 2003; Wang et al., 2013). PXC2 is also known as CANALIZATION-RELATED RECEPTOR-LIKE KINASE (CANAR) and shown to participate in coordination of PIN-FORMED 1 (PIN1) localization during auxin canalization. As a result, *canar* mutants have defects in cotyledon/leaf venation and vascular regeneration after wounding (Hajný et al., 2020). As PXC2 and IRK are both expressed in the root tip, we hypothesized these related LRR-RLKs function redundantly in root development.

Here, we show that PXC2 accumulates in root cell types with a pattern that overlaps with IRK and, like IRK, PXC2 is localized to distinct polar domains or is nonpolar in different cell types. *pxc2* roots show a mild abnormal increase in stele area. However, *irk pxc2* double mutant roots show increases in stele area and endodermal LADs that are more severe than either single mutant. This indicates that polarly localized IRK and PXC2 function redundantly to repress endodermal LADs and stele area in the root meristem. *irk pxc2* double mutant roots also exhibit an abnormal root growth and cotyledon vein patterning phenotype not present in either single mutant, suggesting these receptors are also redundantly required for other aspects of plant development. These double mutant phenotypes can be linked to a role for PXC2 and/or IRK in modulation of auxin-mediated changes in PIN localization. Finally, misexpression of PXC2 can only partially rescue the *irk* root phenotypes, indicating PXC2 is not functionally equivalent to IRK. Thus, PXC2 appears to be partially redundant to IRK with a predominant role for IRK in the root, but these proteins may also have independent functions that could be cell- or tissue-specific. Overall, our results suggest a crucial relationship between the function and polarity of PXC2 and IRK and radial patterning and tropic growth of the Arabidopsis root.

## RESULTS

### PXC2-GFP accumulates in the root tip and is polarly localized

In the root tip, PXC2-GFP expressed under its endogenous promoter (*pPXC2:PXC2:GFP*) shows relatively weak expression, but shows highest accumulation in the LRC, protophloem, and xylem cell types extending shootward from the quiescent center (QC) (Figure 1B, C and S1). PXC2-GFP is also weakly detectable in many other root cell types and in early stage lateral root primordia (Figure 1B and S1A-D). PXC2-GFP is detected at the plasma membrane in root cells, but is also present in the cytoplasm likely in trafficking vesicles. In the LRC, PXC2-GFP is polarly distributed in the plasma membrane at the inner polar domain (Figure 1B, S1B). However, in the phloem and xylem cell types, PXC2-GFP does not show polar accumulation, but when these cells undergo division, PXC2-GFP strongly accumulates in new plasma membranes (Figures 1B and S1B). Due to the very low PXC2-GFP signal, it was difficult to determine in which cell types the reporter accumulated and whether it was polarly localized in cell types other than the LRC.

Given the low accumulation of PXC2-GFP, we used mild plasmolysis combined with cellulase treatment and observed weak PXC2-GFP signal across outer cell types of the root (Figure S1G-I). Additionally, we examined a *pPXC2* transcriptional reporter by expressing endoplasmic reticulum-localized green fluorescent protein (erGFP) with the entire intergenic region (4.7 kilobases) upstream of the *PXC2* start codon. We detected *pPXC2* activity in the lateral root cap and xylem cell types extending shootward from the QC (Figure S1E, F), however its activity was not detected in other cell types (compare Figures 1B, C to S1E, F). Our results are largely consistent with previous examination of *pPXC2* activity via a GUS (β-glucuronidase) reporter (Wang et al., 2013), however, the *pPXC2:GUS* reporter contains just 1.9 kb of the sequence upstream of *PXC2* ((Wang et al., 2013); Dr. Bo Zhang, personal communication). Therefore, differences between these reporters may be due to examination at higher cellular resolution or to differences in presumptive promoter length.

Despite being driven by the same promoter, these *PXC2* reporters show only partially overlapping patterns of expression. We found that expression of *pPXC2:PXC2:GFP* rescues the enhanced *irk pxc2* double mutant back to the *irk* single mutant phenotype (see below), suggesting the accumulation and function of our *pPXC2* driven PXC2-GFP fusion replicates the endogenous situation. The presence of PXC2-GFP in cell types beyond those where *pPXC2* activity is detected may be due insufficient levels of *pPXC2* activity for erGFP detection, whereas PXC2-GFP may be detectable in more cell types due to its accumulation at the plasma membrane. Alternatively, it is possible that PXC2-GFP moves between cells via plasmodesmata. To distinguish between these options and to clearly establish whether PXC2-GFP showed polar accumulation in other cell types, we investigated PXC2-GFP localization using root cell- and/or tissue-specific promoters.

First, we used the *SOMBRERO* promoter (*pSMB,* Willemsen et al, 2008) to drive PXC2-GFP expression specifically in the LRC and confirmed its accumulation at the inner polar domain (Figure 1D). We also expressed PXC2-GFP with *pWEREWOLF* (*pWER*, (Lee and Schiefelbein, 1999) and *pCORTEX2* (*pCO2, (Sabatini et al., 2003; Paquette and Benfey, 2005)* to observe it in the LRC/epidermis and cortex, respectively, and found PXC2-GFP also localized to the inner polar domain in these cell types (Figure 1E, F). In contrast, when PXC2-GFP was expressed from *pSCARECROW* (*pSCR*) (Wysocka-Diller et al., 2000), we observed accumulation at the outer polar domain in the endodermis and the rootward and shootward polar domains in the CEI (Figure 1G). Cell layer-specific expression allowed examination of PXC2-GFP accumulation with less interference from signal in surrounding cells (Alassimone et al., 2010; Campos et al., 2020) and did not reveal any movement of PXC2-GFP between cell types peripheral to the stele. Together, these results indicate that, like IRK, PXC2 is a polarly localized LRR-RLK that accumulates to different polar domains in different cell types (Figure S1J). Overlapping expression domains and identical polar accumulation patterns of IRK and PXC2 in the root are consistent with our hypothesis that these related LRR-RLKs have redundant functions in root development.

### Mutant alleles of *PXC2* have an enlarged stele area

A previous study showed that two *pxc2* alleles had abnormal seed germination in darkness assays (Salk_055351) and were resistant to mannitol (Salk_018730) (ten Hove et al., 2011). These alleles were later found to have defects in cotyledon vein patterning and given the name *canar-1* and *canar-2*, respectively (Hajný et al., 2020). To study PXC2 function in root development, we examined two insertion alleles, SM_3_31635 (*pxc2-3*) and *pxc2-1/canar-1* (Salk_055351) (Figure S2). We confirmed the T-DNA insertion site in *pxc2-3* resides in the first exon and generates a premature stop 10 codons after the insertion site (Figure S2A, not shown). *PXC2* transcript accumulation is reduced in *pxc2-3* (Figure S2B) and based on the position of the insertion, which is upstream of the sequence encoding the transmembrane domain, it is very likely a null allele. The insertion in *pxc2-1/canar-1* is located in the second exon and is downstream of the sequence encoding the transmembrane domain (Figure S2A), suggesting a truncated PXC2 protein could be formed and localized to the plasma membrane similar to truncated versions of IRK (Rodriguez-Furlan et al., 2022). Additionally, we did not find a significant reduction in *PXC2* transcript in *pxc2-1/canar-1* relative to wild type (Figure S2B). We observed that *IRK* expression is somewhat increased in *pxc2-1* seedlings but not in *pxc2-3*, and there is no significant change in *PXC2* transcript levels in *irk-4* (Figure S2C, D). Here, we focus on characterization of *pxc2-3,* as it is likely a complete loss-of-function allele.

No obvious abnormal cellular phenotype was detected in roots of *pxc2-3* seedlings grown on our standard media (not shown). However, when grown on media containing 0.2x Murashige and Skoog (0.2x MS) basal salts, we observed an increase in stele area, but no change in endodermal LADs (Figure 2A, B, E, G). A similar phenotype was also observed in *pxc2-1/canar-1* roots (Figure S3) indicating that both alleles disrupt PXC2 function to restrict stele area. Consistent with our loss-of-function data, inducible overexpression of *PXC2* results in a reduction in stele area (Figure 2F). However, these phenotypic consequences of *PXC2* manipulation are mild and only detectable under particular growth conditions. Thus, PXC2 appears to play a relatively minor role in this process, which may be dependent on environmental conditions and/or partially masked by a redundant gene family member. Unexpectedly, no change in *pxc2-3* cotyledon vein patterning was observed (Figure S4), nor was vein patterning abnormal in *irk-4* (Figure S4E, F). These analyses reveal that *pxc2* mutants exhibit a subset of the root phenotypes observed in *irk-4* (Figure 2F, G), suggesting IRK and PXC2 may have some overlapping or redundant functions in the root, but likely also have independent, cell- or tissue-specific functions.

**Figure 2.**
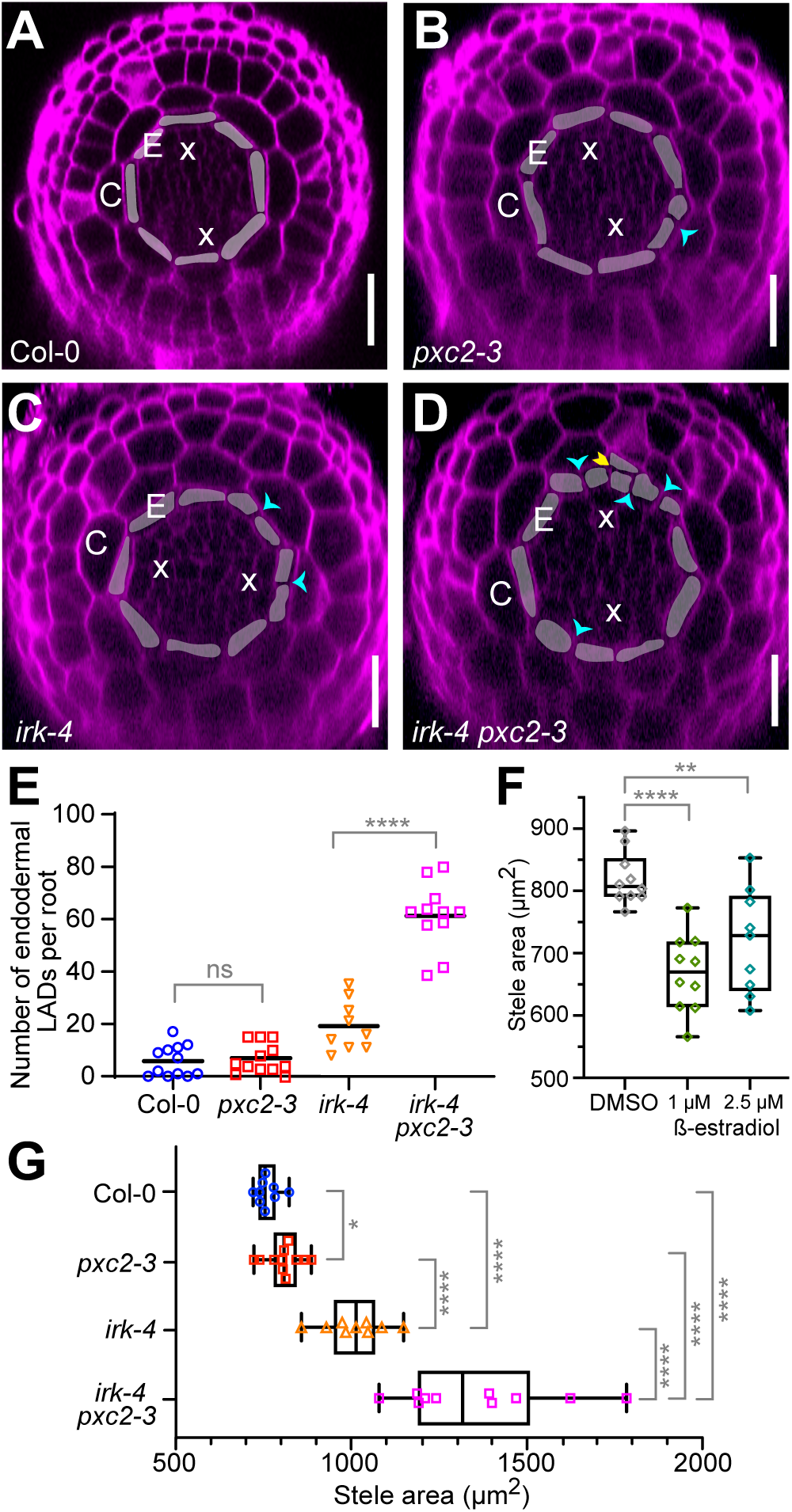
Stele area is impacted in *pxc2-3* and upon PXC2 overexpression and *irk-4 pxc2-3* roots have an enhanced abnormal phenotype. (A-D) Transverse optical sections of Arabidopsis root tips stained with PI (magenta) and endodermal cells shaded, roots at 6 days post stratification (dps) (A, B) and 4 dps (C, D). Cyan arrowheads indicate endodermal LADs and yellow arrowheads indicate periclinal divisions. (E) Scatter plot showing total number of endodermal LADs per root at 6 dps (n = 9-12 roots per genotype). LADs in individual roots represented by colored symbols with the black bar indicating the mean for each data set. (F, G) Box plots showing stele area (F) upon induction of PXC2 overexpression (n = 9-10 roots per treatment) and (G) in various genotypes (n = 9-11 roots per geno-type). Whiskers indicate min/max with interquartile range and median shown with black boxes/lines, respectively, and colored symbols show measurements for individual roots. Data shown are from a single representative biological replicate of ≥3, with similar results. Scale bars: 25 μm. Abbreviations: E - endodermis, C - cortex, x - xylem. Statistics: ns = not significant, p values: * = <0.05, ** = <0.01, **** = <0.0001, Mann-Whitney test.

### *pxc2 irk* double mutants have increased stele area and endodermal longitudinal anticlinal cell division phenotypes

As compared to each single mutant, *irk-4 pxc2-3* roots display an enhanced abnormal phenotype with many excess endodermal LADs and further enlargement of stele area (at 4 dps, Figure 2). The cell division phenotypes observed in *irk-4 pxc2-3* roots become substantially more severe in seedlings at 6 dps (Figure S4A, B) making it difficult to confidently assess the endodermal cell division phenotype, so we deemed cell division quantification reliable only up to 4 dps. To ensure that the phenotypic enhancement observed in *irk-4 pxc2-3* double mutants was due to simultaneous loss-of-function of both genes and not to disruption of any other locus, we expressed *pPXC2:PXC2:GFP* in the *irk-4 pxc2-3* background and observed rescue of the double mutant phenotype to the level of *irk-4* (Figure S4C, D). These results indicate that the increased phenotypic severity of the double mutant is specific to simultaneous loss of IRK and PXC2 function and suggests that these proteins function redundantly to repress endodermal LADs and restrict stele area.

Enlargement of the stele in these mutants may be due to increases in cell size or number. As the stele consists of the pericycle layer and vascular (xylem, phloem, and cambium) cell types (Figure 1A), we first assessed the proportion of the stele area that is made up of pericycle cells and found that in *irk-4* and *irk-4 pxc2-3* the pericycle is proportionally larger (Figure S5). This suggests that in these mutants cells of the pericycle layer are larger in size relative to the vascular cell types. Consistent with this, the number of xylem cells and phloem/cambium cells was not significantly increased in these mutant genotypes (Figure S5A-C). These results indicate stele area increases in these mutants are due to larger cells and not more cell proliferation in the stele.

### *irk pxc2* double mutants exhibit a root growth defect

While investigating the cellular phenotypes in *irk pxc2* roots, we observed a root growth phenotype not present in either single mutant. *irk-4 pxc2-3* roots are shorter and do not grow as consistently and uniformly towards the gravity vector (Figure 3). We quantified several root growth traits and found that when compared to wild type, *irk-4 pxc2-3* mutants show reduced root length and straightness ratio, as determined the distance from hypocotyl to root tip (Dy) divided by total root length (Figure 3A-D). Although in some replicates *irk-4* root length was slightly increased compared to WT that observation was not consistently replicated; thus, we conclude that root length and straightness are not significantly altered in *pxc2-3* and *irk-4* single mutants (Figure S5D, E). However, in contrast to WT roots, which exhibit a weak right-slanting (Dx) growth habit (Grabov et al., 2005; Arribas-Hernández et al., 2020), all mutant genotypes show a left-slanting growth habit (Figures 3A, B, F-H). Finally, upon reorientation of the seedlings with respect to the gravity vector, *irk-4 pxc2-3* mutant roots show an attenuated gravitropic response compared to wild type, while the single mutants showed a wild-type response (Figures 3E and S5F). The abnormal growth and gravi-stimulation response are unique to *irk pxc2* double mutants but do not strongly impact overall growth and maturation of these plants on soil (not shown). Additionally, while lateral root capacity in *pxc2-3* single mutants is slightly increased, it is slightly reduced in *irk-4* and the double mutants (Figure S5G), suggesting no major role for IRK and/or PXC2 in specifying sites competent for lateral root formation. Overall, these results extend the functional requirement for IRK and PXC2 to include a redundant role for them in root growth.

**Figure 3.**
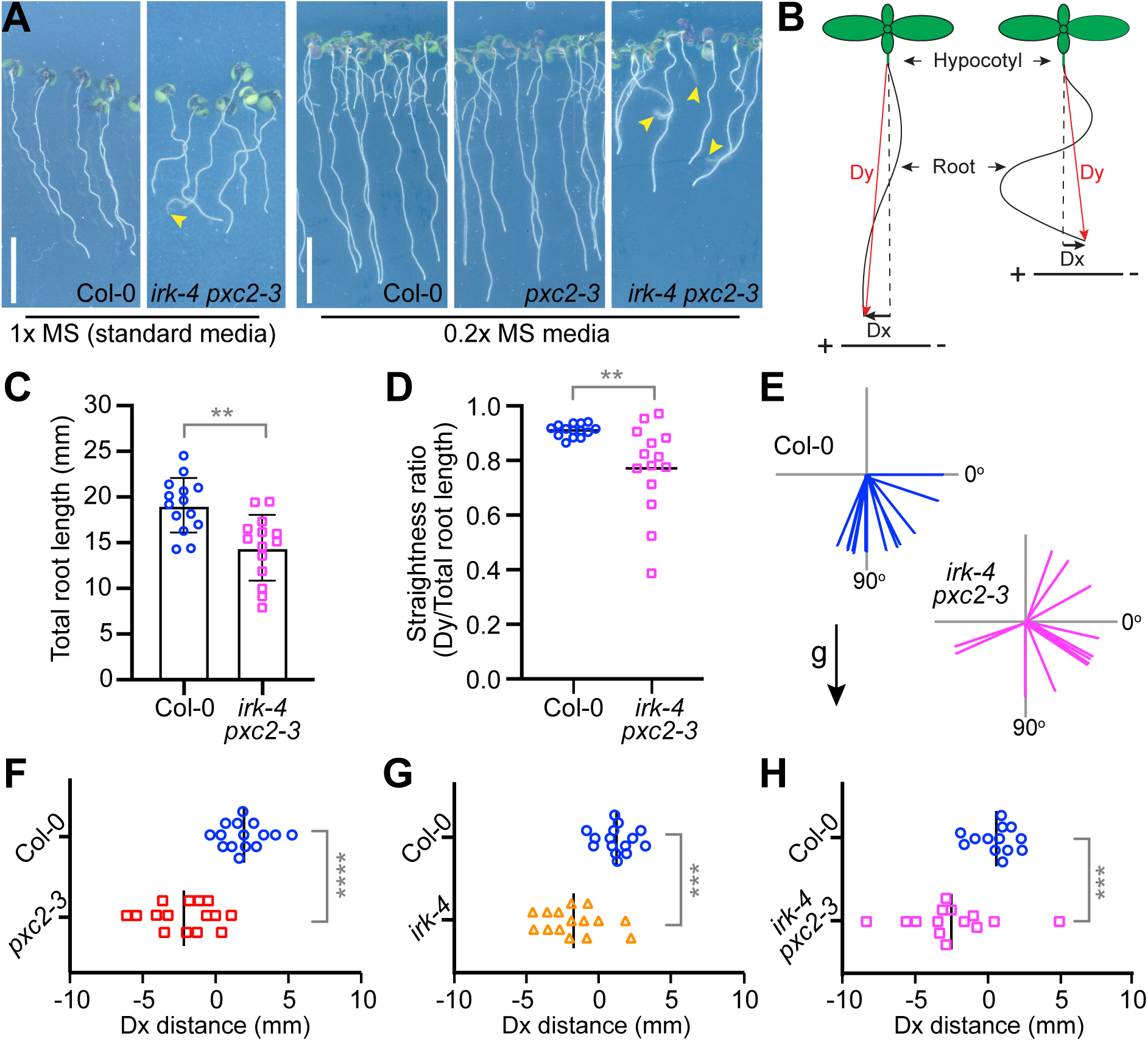
*irk pxc2* roots exhibit abnormal growth phenotypes. (A) Seedlings at 7 dps grown on 1x MS (our standard growth medium) and 0.2x MS plates sealed with micropo-re tape. Scale bars = 10 mm, yellow arrowheads indicate root tips growing off the media surface. (B) Schematics of seedlings with different root growth phenotypes and measurements used to calculate straightness ratio and Dx distance. (C-H) Quantification of root phenotypes of plants grown on 1x MS. (C-E) Comparison of Col-0 and *irk-4 pxc2-3* roots at 7 dps (C) Average total root length (error bars = standard deviation) and (D) straight-ness ratio (Dy/total root length, n =14 per genotype). (E) Angle of Col-0 (n = 14) and *irk-4 pxc2-3* (n = 13) root tips 8h after gravistimulation (plates turned 90°). Lines indicate angles of individual root tips with respect to the new gravity vector indicated by ‘g’ and black arrow (previous gravity vector aligned to 0°). (F-H) Graphs of Dx distance in wild type compared to (F) *pxc2-3* (n = 15 per genotype), (G) *irk-4* (n = 15 per genotype), (H) *irk-4 pxc2-3* (n = 13 per genotype). (D, F-H) Scatter plots show data for individual roots indicated by colored symbols with the black bar indicating the mean for each data set. All data shown is from a single biological replicate of ≥3. Statistics: p-value = ** <0.005, *** <0.001, **** <0.0001, Mann-Whitney test.

### IRK and PXC2 modulate auxin feedback on PIN1 localization and trafficking

The abnormal root growth phenotype of *irk pxc2* double mutants may be attributed to the presence of ground tissue cell files that are non-uniform in cell number and shape, as directional growth requires the coordinated elongation of neighboring cell types (Vaahtera et al., 2019; Sablowski, 2016). Alternatively, root gravitropism requires the (re)polarization of PINs to achieve differential auxin distribution and cell elongation (Kleine-Vehn et al., 2010; Friml et al., 2002) and PXC2/CANAR binds to and coordinates the polarization of PIN1, an auxin efflux carrier (Hajný et al., 2020). We found that *irk-4 pxc2-3* often have cotyledon vein patterns with more than four vascular loops compared to wild type or either single mutant (Figure S4E, F). These phenotypes suggest PXC2 and IRK may be redundantly required for normal polar auxin transport. Unexpectedly, we also found that *irk-4* and *irk-4 pxc2-3* endodermal cells do not relocalize PIN1 to the inner lateral PM domain in response to exogenous auxin (naphthaleneacetic acid (NAA)), as WT and *pxc2-3* do (Figure S6). Together these results suggest auxin-induced PIN1 repolarization in the endodermis requires IRK.

Although the precise mechanisms of auxin-induced PIN repolarization remain largely elusive, the canonical TIR1/AFB and cell surface ABP1-TMK pathways are required, as well as endocytic trafficking (Mazur et al., 2020b; Friml et al., 2022; Mazur et al., 2020a). In Arabidopsis, Brefeldin A (BFA) disrupts endocytic trafficking by inhibiting protein secretion and recycling, which triggers “BFA-body” formation (Kleine-Vehn et al., 2008; Han et al., 2021). Treatment with NAA, inhibits BFA-body formation, suggesting exogenous auxin influences PIN endocytic trafficking (Narasimhan et al., 2021), implicating auxin feedback on PIN polarity (Wabnik et al., 2010). To further examine a role for IRK and PXC2 in auxin-induced PIN1 endocytic trafficking, we treated roots with BFA and NAA and quantified BFA-body size and density. In the pericycle of wild type and the single mutants, the number and size of BFA-bodies formed under control and NAA treatments were similar (Figure S6C-E). However, in control conditions, *irk-4 pxc2-3* roots had smaller BFA-bodies with an average size of 1.88 μm^2^, compared to WT bodies at 2.42 μm^2^. Despite this, all genotypes showed similar BFA-body size upon NAA treatment (Figure S6C-D). NAA treatment also decreased the density of BFA-bodies in all genotypes, except *irk-4 pxc2-3* where BFA-body density was slightly higher (Figure S6E). These results suggest that IRK and PXC2 have partially redundant functions in PIN1 trafficking and its response to exogenous auxin, potentially linking these receptors to auxin-mediated PIN repolarization.

### PXC2 is not functionally equivalent to IRK

PXC2 and IRK share 57.1% identity across their entire amino acid sequence and 65.1% identity within their intracellular domains (Figure S7). Additionally, they have overlapping expression and polar accumulation patterns. *pxc2* roots exhibit a subset of the abnormal phenotypes observed in *irk* and the *irk pxc2* double mutant root phenotype is more severe, which suggests functional redundancy. Some related, functionally redundant LRR-RLKs are also functionally equivalent upon misexpression. For example, ERECTA (ER) is 62-63% identical to its functional paralogs ERECTA-LIKE 1 (ERL1) and ERL2 and they are functionally equivalent to ER (Shpak et al., 2004). This was also observed for BRI1 and closely related BRI1-LIKE 1 (BRL1) and BRL3, which are 49% identical to BRI1 (Caño-Delgado et al., 2004). As PXC2 and IRK have comparable amino acid identity, we explored whether PXC2 could functionally replace IRK in the endodermis.

PXC2 and IRK are both expressed and polarly accumulate in the endodermis, albeit IRK is expressed there at a higher level. *pSCR-*driven misexpression of IRK-GFP was shown to be sufficient to rescue the increased endodermal LADs and largely rescue stele area in *irk-4* (Campos et al., 2020). We assessed whether *pSCR* driven PXC2-GFP could similarly rescue *irk-4* (Figure 4). In *irk-4* seedlings that were homozygous for *pSCR:PXC2:GFP* endodermal LADs were slightly increased but not statistically different from wild type, while the stele area of these roots was significantly reduced compared to *irk-4*, but not fully rescued (Figure 4A, B). However, in older *irk-4* seedlings (6 dps) segregating for *pSCR:PXC2:GFP*, we observe only partial rescue of both the endodermal LAD and stele area phenotypes (Figure 4C, D). These results indicate that endodermal-expressed *PXC2* cannot fully compensate for the loss of IRK in that cell type. This shows that PXC2 is not functionally equivalent to IRK and suggests these proteins have some distinct functions, which may be cell type-specific.

**Figure 4.**
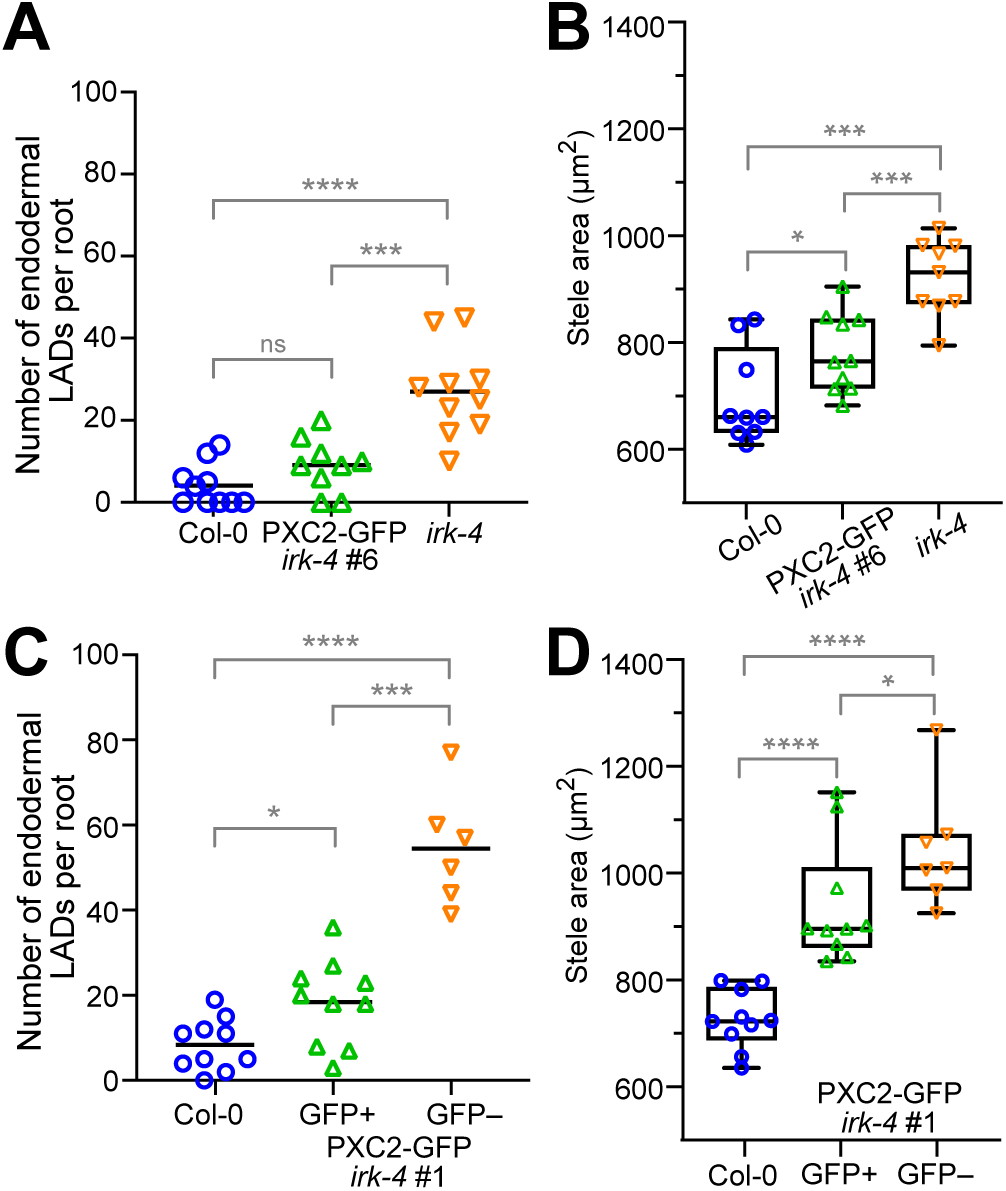
Misexpressed PXC2 partially rescues abnormal *irk* phenotypes. Quantification of the total number of endodermal LADs and stele area in (A, B) Col-0, *irk-4 pSCR:PXC2:GFP*, and *irk-4* roots (at 4 dps, 9-10 roots per genotype) and (C, D) Col-0 and *irk-4* segregating for *pSCR:PXC2:GFP* (at 6 dps, Col-0 n = 10, GFP positive n = 10, GFP negative, n = 7). All data shown are from a single, representative biological replicate of ≥3. Scatter plots (A, C): Total number of endodermal LADs per root indicated by colored symbols and black bar indicating the mean for each data set. Box plots (B, D): whiskers indicate min/max with interquartile range and median shown in black boxes/lines, respectively, and colored symbols indicating measurements for individual roots. Abbreviations: ns = not significant, p-values: * = <0.05, *** ≤ 0.001; **** ≤ 0.0001, Mann-Whitney test.

## DISCUSSION

With the characterization of PXC2, we have identified another LRR-RLK putatively involved in polarized cell-cell communication and another player in root ground tissue cell division and patterning. Examination of *pxc2* mutant roots revealed increased stele area compared to WT. This phenotype is mild and observed only when seedlings were grown on media containing a lower concentration of MS salts than our standard growth media. This suggests the consequences of loss of PXC2 function are exacerbated by lower nutrient conditions or more rapid root growth. Because IRK and PXC2 are closely related and the *irk-4* root phenotypes are enhanced in *irk-4 pxc2-3* double mutants, we propose that PXC2 and IRK act redundantly to restrict stele area and repress endodermal LADs, such that there are typically eight endodermal cells around the stele (Figure 5). Given the mild abnormal *pxc2* phenotype, our data are consistent with a predominant role for IRK in these processes. The enlarged stele of *irk* and *pxc2* single mutants, suggests both proteins are required to restrict stele area. It is possible that IRK and PXC2 have independent functions in this process, however, given their phylogenetic relationship (Shiu and Bleecker, 2001), we propose these proteins function redundantly and that a lower dosage of the gene products leads to weaker single mutant phenotypes (Figure 5, center). Because *pxc2* single mutants do not have excess endodermal LADs, but this phenotype is dramatically enhanced in *irk pxc2* roots compared to *irk* alone, we propose that IRK is fully redundant with PXC2 to prevent these endodermal divisions.

**Figure 5.**
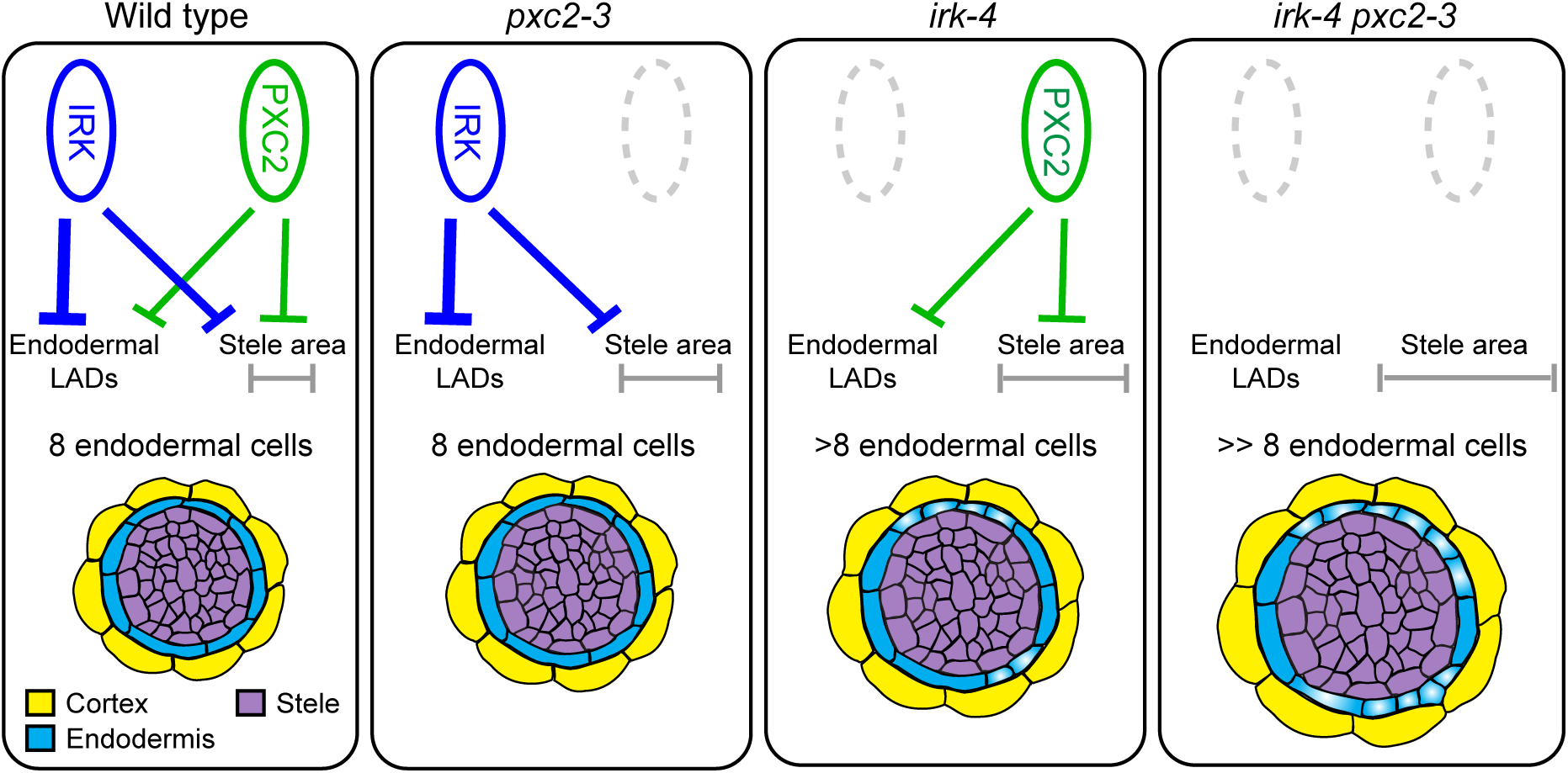
Summary of IRK and PXC2 root functions. Diagrams illustrating the roles of IRK and PXC2 with regard to repression of endodermal LADs and stele area (grey bracket). Weight of the lines indicates proposed contribution of IRK (blue) and PXC2 (green) with the phenotypic outcomes listed. (lower) Schematic representations of transverse sections of a portion of the root showing phenotypes observed in endodermal cells and stele in Col-0, *pxc2-3*, *irk-4*, and *irk-4 pxc2-3*. Highlighted endodermal cells indicate the occurrence of LADs; therefore, more endodermal cells in the root’s radial axis.

Our analyses indicate a role for IRK and PXC2 beyond repression of endodermal LADs and stele area. Each single mutant shows left-slanted root growth along with increased stele area suggesting these phenotypes may be related. These phenotypes are both more severe in the double mutant, which also exhibits an agravitropic root growth phenotype and cotyledon vein patterning defect, not observed in either single mutant. This suggests a broader role for these proteins in plant development that appears linked to modulation of polar auxin transport. Auxin-based feedback on PIN1 polarity and trafficking is considered an essential prerequisite of auxin canalization and (re)direction of polar auxin transport and PXC2 interacts with PIN1 (Hajný et al., 2022; Wabnik et al., 2010). We find that PIN1 repolarization is impacted in *irk-4* and *irk-4 pxc2-3*, however this phenotype is not more severe in the double mutant suggesting IRK alone participates in this process. Yet in the presence of BFA, only the double mutant shows a decrease in PIN1-BFA-body size and, upon BFA+NAA treatment, only the double mutant has more BFA-bodies per cell. This suggests that IRK and PXC2 have partially redundant functions in PIN trafficking and auxin feedback on PIN polarity, which can explain the agravitropic root growth of the double mutant.

Yet these results were a bit unexpected as the *pxc2-1/canar-1* allele shows reduced PIN1 reorientation in response to NAA and reduced BFA-bodies per cell upon BFA treatment in roots (Hajný et al., 2020). The T-DNA insertion responsible for the *pxc2-1/canar-1* and *pxc2-2/canar-2* alleles are located within the 2nd exon, which encodes the kinase domain, suggesting that a truncated version of PXC2 could be produced in these plants. Our results with IRK indicate that a kinase-deletion version remains polarly localized and is partially functional (Rodriguez-Furlan et al., 2022). Kinase truncations in transmembrane receptor kinases can have a range of molecular and phenotypic consequences. We propose there is a unique feature of *pxc2-1/canar-1* allele that results in some distinct phenotypic consequences compared to the null allele. Given the similarity between IRK and PXC2 it might be predicted that, like PXC2, IRK has similar interactions with PINs. Further investigation into how these receptors participate in the mechanisms underlying polar auxin transport and whether this is related to their functions to repress endodermal LADs and stele area will be an important avenue of future work.

Under its own promoter, we show that PXC2-GFP is weakly expressed throughout the root with stronger expression in the LRC, where it accumulates at the inner polar domain, and in the protophloem and xylem cell types, where it appears nonpolar. Similar to IRK, misexpression of PXC2-GFP reveals polar accumulation in various cell types. Their accumulation at distinct polar domains in different cell types suggests localization of IRK and PXC2 is informed by adjacent cells and not organ-level or global polarity cues as proposed for polarly localized nutrient transporters (Alassimone et al., 2010; Takano et al., 2010; Barberon et al., 2014) and the SOSEKI proteins (Yoshida et al., 2019), respectively. We have shown that IRK polar accumulation occurs via targeted secretion with sorting determinants unexpectedly residing in its extracellular domain (Rodriguez-Furlan et al., 2022). Given their amino acid similarities, we predict that IRK and PXC2 share common polarization mechanisms.

Although IRK and PXC2 are expressed in the stele cell types, their expression patterns only partially overlap. IRK-GFP is detected in the pericycle and *pIRK* is active at the phloem poles, but excluded from the xylem (Campos et al., 2020), whereas PXC2-GFP is detected in the pericycle, protophloem and xylem cell types. Given that single mutants of IRK and PXC2 have distinguishable root phenotypes and the double mutant root phenotype is an enhancement of that observed in the single mutants, these RLKs appear to have some redundant and unique roles in the plant. These functional differences might be attributable to differences in the endogenous protein expression and accumulation in each root cell type, such that the primary site of PXC2 function is in the stele, while that of IRK is in the endodermis.

Although it is more broadly expressed, endodermal-expressed IRK rescued endodermal LADs and largely rescued stele area, suggesting that restriction of the endodermis to eight cells around the stele was sufficient to constrain stele area in *irk-4*. Thus, we proposed that increased stele area in *irk-4* was a secondary defect resulting from increased endodermal cell numbers in the radial axis due to excess LADs (Campos et al., 2020). Consistent with this, root vascular patterning and symmetry was recently connected to compressive stress on vascular cell types, likely from the surrounding cell layers (endodermis and pericycle) (Fujiwara et al., 2023). However, our data suggest there may be more complexity in the relationship between endodermal LADs and stele area increases and the roles of IRK and PXC2 in repressing them.

Here, we find stele area increases in *pxc2* roots without a coincident increase in endodermal LADs. If an increase in stele size precedes endodermal LADs, perhaps in *pxc2* some stele size threshold that would trigger abnormal endodermal LADs has not been reached, whereas stele enlargement in *irk-4* reaches that threshold earlier. Consistent with this, *irk-4* roots expressing *pSCR:PXC2:GFP* show increased stele area before excess endodermal LADs, supporting the idea that stele area increase over time may lead to endodermal LADs. Thus, it is possible that in *irk* and *irk pxc2* roots, dysregulation or uncoupling of communication between the endodermis and stele leads to the observed phenotypes, pointing towards mechanical feedback between these cell types in the root’s radial axis. Specifically, excess endodermal LADs would broaden the root’s radial axis, potentially reducing compressive stress allowing stele area to increase. On the other hand, if the stele was enlarged, endodermal LADs may occur to relieve the increased compressive stress on the internal cells. It is noteworthy that in genotypes with the most numerous endodermal LADs (*irk-4 and irk-4 pxc2-3*) the neighboring cell layer, the pericycle, takes up a significantly larger proportion of the enlarged stele. Further exploration of the cell type-specific functions of IRK and PXC2 during root development will allow deeper functional insights into these proteins, and importantly, can serve as novel tools to dissect the interaction between endogenous genetic control and exogenous mechanical feedback during developmental patterning. Given our collective results, we hypothesize a crucial relationship exists between the polarity and function of PXC2 and IRK in radial patterning and tropic growth of the Arabidopsis root.

## EXPERIMENTAL PROCEDURES

### Plant Materials and Growth Conditions

Seeds were surface sterilized with chlorine gas, then plated on media (pH 5.7) containing 1% (BD Difco™) Agar and 0.5g/L MES (EMD), supplemented with 1% sucrose w/v and 1x Murashige and Skoog (MS, Caisson labs) basal salts (our standard growth medium) or 0.2x MS salts (as noted). Media containing 0.2x MS salts increases root growth rate due to lower levels of nitrogen and potassium but does not alter root development. Plates were sealed with parafilm or 3M micropore tape as indicated below. Seeds were stratified on plates in the dark at 4°C for 24-72 hours and then placed vertically in a Percival growth chamber, with a 16h light/8h dark cycle at a constant temperature of 22°C for 4-7 days post-stratification (dps), unless otherwise noted. PXC2 overexpression (*35S:XVE:PXC2:3HA* in Col-0, details below) was induced with 1 µM and 2.5 µM β-Estradiol (solubilized in DMSO, primary stock at 10 mM) on 0.2x MS plates sealed with parafilm until imaging at 6 dps.

### Vector Construction and Plant Transformation

Transcriptional and translational reporters were constructed by standard molecular biology methods and utilizing Invitrogen Multisite Gateway® technology (Carlsbad, USA). The 4.7 kb region upstream of the *PXC2* (At5g01890) start codon was amplified from Col-0 genomic DNA and recombined into the Invitrogen pENTR^TM^ 5’-TOPO® TA vector as the promoter of *PXC2*. For the transcriptional reporter, *pPXC2* drove endoplasmic reticulum-localized green fluorescent protein (erGFP) as previously described (Van Norman et al., 2014). For translational fusions, the genomic fragment encoding *PXC2* from the ATG up to, but excluding the stop codon (including introns, 3.0 kb), was amplified from Col-0 genomic DNA and recombined into the Invitrogen pENTR^TM^ DIRECTIONAL TOPO® (pENTR-D-TOPO) vector and fused to a C terminal GFP tag as previously described (Van Norman et al., 2014). Using overlapping PCR, the genomic fragment encoding *PXC2* without its stop codon was C-terminally fused to an 3xHA tag via linker (5x glycine). The PXC2-3HA fragment was recombined into the pDONR221 gateway entry vector via BP reaction. This entry vector was subsequently recombined into pMDC7 destination vector with XVE inducible promoter via LR reaction. Specific primers for *PXC2* cloning are listed in Table S1.

Cell type- or layer-specific promoters (*pSCR, pCO2,* and *pWER)* as previously described (Campos et al., 2020; Lee et al., 2006), and *pSMB* as described (Bennett et al., 2010) were used to drive PXC2-GFP. The various Gateway compatible fragments were recombined together with the dpGreen-BarT (Lee et al., 2006) or dpGreen-NorfT (Norflurazon resistant) destination vectors. The dpGreenNorfT was generated by combining the backbone of dpGreenBarT with the *p35S::tpCRT1* and terminator insert from *pGII0125*. Within the target region of the dpGreenBarT, one AclI site was mutated with the QuickChangeXL kit (Stratagene) and primers as in Table S1. Plasmids were amplified in ccdB-resistant *E. coli* and plasmids prepped with a Bio Basic Plasmid DNA Miniprep kit. 34uL of the modified dpGreenBarT and unmodified *pGII0125* were digested with 1ul each FspI and AclI in CutSmart buffer (NEB) for 1hr at 37°C. Digests were subjected to gel electrophoresis on a 1% agarose gel. The 5866bp fragment from the dpGreenBarT and 2592bp fragment from the *pGlIII0125* were extracted with a Qiagen MinElute Gel Extraction kit. The fragments were then ligated at 1:1 volumetric ratio (20ng vector; 8.8ng insert) using T4 DNA ligase incubated at 16°C overnight before transformation into ccdB-resistant *E. coli*.

Expression vectors were then transformed into Agrobacterium strain GV3101 (Koncz et al., 1992) and then into Col-0 plants by the floral dip method (Clough and Bent, 1998). Transformants were identified using standard methods. For each reporter, T2 lines with a 3:1 ratio of resistant:sensitive seedlings, indicating the transgene is inherited as a single locus, were selected for propagation. These T2 plants were allowed to self and among the subsequent T3 progeny, those with 100% resistant seedlings, indicating that the transgene was homozygous, were used in further analyses. For each reporter, at least three independent lines with the similar relative expression levels and localization patterns were selected for imaging by confocal microscopy.

### Confocal microscopy and analysis of fluorescent reporters

Roots were stained with ∼10 μM propidium iodide (PI) solubilized in water for 1-2 min and visualized via laser scanning confocal microscopy on a Leica SP8 upright microscope housed in the Van Norman lab. Root meristems were visualized in the median longitudinal, transverse planes, or as Z-stacks. Fluorescence signals were visualized as follows: GFP/YFP (excitation 488 nm, emission 492-530 nm) and PI (excitation 536 nm, emission 585-660 nm). All confocal images are either median longitudinal optical sections, transverse optical sections, or part of a Z-stack acquired in the root meristematic or elongation zone. Images of roots expressing *PXC2* transcriptional or translational fusions were taken at 4-7 dps with no observable differences due to age. Roots misexpressing PXC2-FP using cell type-specific promoters were imaged at 5 dps.

To assess PXC-GFP accumulation in partially plasmolyzed cells, *pPXC2:PXC2-GFP* expressing seedlings (5-7 dps) were placed in 1x MS liquid media and treated with 20 μM FM4-64 for 5 min, rinsed with 1x MS liquid media, and placed in 0.4 M D-Mannitol, 20 mM MES monohydrate, 20 mM KCl, 10 mM CaCl_2_, and 3% cellulase (pH adjusted to 5.7 with 1 M Tris-HCl at pH 7.5) for the indicated times. Fluorescent signals were visualized using a 488 nm laser to excite GFP and FM4-64, with GFP emission collected at 492–530 nm and FM4-64 emission at 600–660 nm. Images were acquired by line sequential scanning to avoid signal bleed through. Image processing was performed using the LAS X software and ImageJ software (<u>imagej.nih.gov/ij/</u>).

### Endodermal LAD quantification and stele area measurement

All images of roots for analysis of endodermal LADs and stele area were taken as Z stacks at 512×512 resolution with 1 µm between scans, speed setting at 600 with a line average of six. Z-compensation was used by increasing 588 nm laser intensity while focusing through the Z plane, from 2% up to 40% (depending on staining) in order to visualize cell layers adjacent to the slide (in the radial axis).

Total number of endodermal LADs were determined by counting the number of endodermal cells per ring of cells in the transverse plane starting from just above the QC. Because cells of the cortex and endodermis are formed as a pair through periclinal division of a single initial cell, endodermal LADs typically result in two or more small endodermal cells adjacent to a single cortex cell. For each individual root, 15 endodermal rings were scored for the occurrence of LADs, the presence of an additional longitudinal anticlinal wall within a ring of endodermal cells was scored as an LAD and then summed. If any of the first 15 endodermal cells were damaged (PI infiltration) the root was not used in the analysis.

Stele area was measured in the transverse section located 60μm above the QC using ImageJ software. The inner cell wall of the endodermis was traced and used to calculate the area of the region in ImageJ. If the stele was damaged or staining was too weak to accurately determine the endodermal border then that root was excluded from the analysis.

### Phenotypic characterization of *pxc2* and *irk-4 pxc2* mutants

*pxc2-3/canar-3* (SM_3_31635) and *pxc2-1/canar-1* (Salk_055351) seeds were obtained through the Arabidopsis Biological Research Center and genotyped using primers listed in Table S1 to identify homozygous mutants. Age matched seeds from plants grown for three generations in the lab were used in these studies. Each mutant was plated with Col-0 on individual plates. Seeds were sown on plates containing 0.2x MS media and sealed with paraflim. At 6 dps, Z-stacks were obtained for *pxc2* mutants and Col-0 root tips (n = 10-15 per genotype) for quantification of endodermal LADs and stele area. No additional root cellular morphology phenotype was observed for either *pxc2* mutant allele.

Standard genetic crosses between *pxc2-3* and *irk-4* plants were completed to generate the *irk-4 pxc2-3* double mutant. F2 plants were genotyped to identify genotypes of interest and then confirmed in subsequent generations. Double mutant seeds collected from genotyped F3s were used in these studies. *irk-4 pxc2-3* mutants were plated with *irk-4* as the controls on plates containing 1x MS media and sealed with parafilm. At 4 dps, Z-stacks were obtained for *irk-4 pxc2-3* and *irk-4* (n = 10-15 per genotype) for quantification of endodermal LADs and stele area. Additional imaging was conducted at 6 dps, however, quantification of stele area and endodermal LAD was not performed due to severe root tissue patterning defects.

For the root length, straightness ratio, and gravi-stimulation analyses, Col-0, *pxc2-3*, *irk 4*, and *irk-4 pxc2-3* seeds were plated on our standard media and sealed with micropore tape and plates were scanned (EPSON V600) at 7 dps. After scanning, the plates were turned 90° and placed back into the growth chamber for 8 hours, then scanned again. From the first scans, total root length was measured by tracing individual roots from the base of the hypocotyl to the root tip using the segmented line feature in ImageJ. Then, a straight line was drawn from the base of the hypocotyl to the root tip and measured as Dy (Figure 4A). The straightness ratio is calculated by dividing Dy by total root length where a ratio of 1.0 indicates root growth always parallel with the gravity vector. To assess Dx (Figure 4A), a straight line was first drawn directly downward from the base of the hypocotyl to the region parallel to the root tip. Then, a straight line was drawn from this point to the root tip, resulting in the Dx measurement, with a positive value assigned to the rightward direction of root drift as observed in Col-0. From the second scans (8 hours after gravistimulation), the angle between 1mm of the root tip and the new gravity vector were measured in ImageJ to assess the root’s response following gravistimulation.

For cotyledon vein patterning analyses, 10-day old seedlings were imaged after growth on 0.5x MS medium for 11 dps (sealed with tape). Shoot tissue, including cotyledons of each genotype was harvested and then subjected to 30-min fixation using ethanol:acetic acid (3:1) fixation (three times). The fixed samples were incubated with 75% ethanol for washing and clearing, and then imaged on an Olympus BX53. Vein patterns with 4 fully connected loops or base loops open at the bottom were considered “4 loops, normal”, while those with any opening in the apical loops or at the middle/top position in the base loops were considered “4 loops abnormal”.

### PIN1 localization studies

Whole-mount *in situ* immunolocalization was conducted to observe PIN1 lateralization and BFA-body formation as described previously (Sauer et al., 2006; Han et al., 2021). For PIN1 lateralization in roots, 4-day old plants grown on 0.5x MS plates (sealed with tape) were transferred to liquid 0.5x MS-medium containing 10 μM NAA or DMSO and incubated for 4 hours before fixation. A 20 mM NAA stock was diluted in DMSO and DMSO was used as the control treatment. For BFA-body observations, 4-day old plants grown on MS plates were incubated in liquid MS medium containing 50 μM BFA and 10 μM NAA or DMSO for 2 hours before fixation. The *in situ* immunolocalization used these antibodies: anti-PIN1 (1:1000) and secondary goat anti-rabbit antibody coupled Cy3 (Sigma-Aldrich, 1:600). Confocal imaging was carried out on a Zeiss LSM 800 microscope (Cy3, Excitation 548 nm and Emission 561 nm). PIN1 lateralization and BFA-bodies were measured using Fiji (v10), Lookup Tables, with 16 color (for PIN1 lateralization assay) and Magenta Hot (for BFA treatment) intensity scales used to indicate PIN1 accumulation in roots.

### PXC2-GFP rescue of *irk-4*

*irk-4* plants homozygous for *pSCR:PXC2:GFP*, Col-0, and *irk-4* mutants were plated on 1x MS media and imaged at 4 dps as previously described. *irk-4* mutants segregating for *pSCR:PXC2:GFP* and Col-0 were plated together on media previously described and imaged at 6 dps. Two independent transgenic lines (#4 and #6) were crossed with *irk-4* for analysis at both 4 and 6 dps with similar results.

### Figure construction

Confocal images for stele area measurement and endodermal LAD quantification were examined in ImageJ (http://imagej.nih.gov/ij) (Schneider et al., 2012). For use in figures, raw confocal images were converted to .TIF format using Leica software (LASX) and were assembled in Adobe Photoshop. Statistical analysis and graphical representation of all data in this publication was done using PRISM8 (GraphPad Software, https://www.graphpad.com/, San Diego, USA) or OriginPro 2022 (OriginLab, https://www.originlab.com/origin). Type of graphs and statistical analysis used are listed in figure legends. Schematics were created in Illustrator and then figures containing images, graphs, and schematics were assembled in Adobe Illustrator.

### qRT-PCR analysis

Total RNA was isolated from seedlings at 7dps of three independent biological replicates for each Col-0 (wild type), SM_3_31635 (*pxc2-3/canar-3*) and Salk_055351 (*pxc2-1/canar-1*) using Qiagen RNeasy Plant Mini Kit. Seeds of all three genotypes were sown together on plates containing 1x MS media, sealed with parafilm, and stratified in the dark at 4°C for 3 overnights. Following total RNA extraction, samples were examined for concentration and purity and stored at −80°C prior to cDNA synthesis. For cDNA synthesis (RevertAid First Strand cDNA Synthesis Kit,Thermo Scientific), 1μg of total RNA was used to normalize for varying RNA concentrations within biological samples, and the Oligo(dT)18 primer was used to generate cDNA. To examine *PXC2* expression in *irk-4*, previously obtained RNA samples (Campos et al., 2020) were used. qRT-PCR reactions were done using IQ SYBR Green Supermix (BioRad) and analysis was performed on the CFX Connect Real-Time System housed in the Integrative Institute of Genome Biology Genomics Core facility at UC-Riverside. The reaction conditions for all primer pairs were: 95°C for 3 min, followed by 40 cycles of 95°C for 10s and 60°C for 30s. Primer pair efficiency was calculated for each reaction using standard curve data. For each genotype and biological replicate, three technical replicates were performed, and all transcript levels were normalized to *PHOSPHATASE 2A* (*PP2A*) (Czechowski et al., 2005). Data analysis was done using Bio-Rad CFX Maestro 1.1 (version 4.1.2433.1219).

### Gene structure and amino acid sequence comparisons

DNA sequence of *PXC2* including 5’ and 3’ UTR was copied into the web-based http://wormweb.org/exonintron tool and annotated with the positions of the T-DNA insertions based on DNA sequencing for *pxc2-3* and using publicly available data from The Arabidopsis Information Resource (TAIR, www.arabidopsis.org) for *pxc2-1/canar-1* and *pxc2-2/canar-2*. Sequence identity and similarity of the intracellular regions of IRK and PXC2 was done using BLAST local alignment tool (https://blast.ncbi.nlm.nih.gov/Blast.cgi) protein blast. Visual representation of intracellular regions alignment of IRK and PXC2 were done by pasting FASTA sequences into T-Coffee (http://tcoffee.crg.cat/apps/tcoffee/index.html) multiple sequence alignment tool and sequences were downloaded as “fasta_aln„ file. This alignment was then formatted using Boxshade (https://embnet.vital-it.ch/software/BOX_form.html), using “RTF_new„ as the output format and downloading the output file as a Word document. Prediction of the transmembrane domain region was done using TMHMM Server 2.0 (https://services.healthtech.dtu.dk/services/TMHMM-2.0/). Percent identity for PXC2 and IRK and of ERL1 and ERL2 to ER and BRL1 and BRL3 to BRI1 were obtained using the web based Clustal Omega tool (https://www.ebi.ac.uk/Tools/msa/clustalo/).

## ACKNOWLEDGMENTS

We thank Roya Campos for assistance in the construction of several PXC2-GFP misexpression vectors and Dr. Dawn Nagel, Dr. Patricia Springer, Roya Campos, and Jessica N. Toth for discussions of the project and feedback on the manuscript while it was in preparation. We also thank Dr. Erin Sparks (University of Delaware) for providing the NorfT Gateway® compatible destination vector. We appreciate access to and assistance from the Institute of Integrative Genome Biology Genomics Core Facility (UC, Riverside) for qRT-PCR experiments. Research conducted in the Friml lab was supported by Austrian Science Fund (FWF) (#I3630-775 B25) and the Scientific Service Units of ISTA through resources provided by the Imaging & Optics Facility and the Lab Support Facility. The work of Jakub Hajný was supported by a long-term EMBO fellowship (ALTF217-2021). Work in the Van Norman lab was supported funds awarded to JMVN, specifically by Initial Compliment (IC) funds from the University of California at Riverside, USDA-NIFA-CA-R-BPS-5156-H, and by an NSF CAREER award (#1751385).

## SUPPORTING INFORMATION

**Figure S1:** P*X*C2 promoter activity and PXC2-GFP accumulation in the primary and lateral root primordia.

**Figure S2:** P*X*C2 gene structure and expression analyses.

**Figure S3:** p*x*c2*-3* and *pxc2-1/canar-1* roots have similar abnormal root phenotypes.

**Figure S4:** *irk-4 pxc2-3* root and cotyledon vein patterning phenotypes with root phenotype rescued back to *irk-4* by expression of *pPXC2:PXC2:GFP*.

**Figure S5:** Stele cell and root growth phenotypes in *pxc2-3* and *irk-4* single and double mutants.

**Figure S6:** IRK and PXC2 participate in auxin-mediated feedback on PIN1 accumulation and trafficking in roots.

**Figure S7.** Alignment of IRK and PXC2 intracellular domains.

**Table S1:** List of primers used in this study.

## AUTHOR CONTRIBUTIONS

Conceptualization: J.M.V.N.; Methodology and Investigation: PXC2 localization: J.M.V.N., J.G., I.K.R, and C.R.F.; root phenotyping: J.M.V.N., J.G., I.K.R.; PXC2 inducible overexpression: J.H. and I.K.R.; Cotyledon vein patterning: J.H., Z.G., and J.F.; PIN1 localization and BFA-body formation: Z.G. and J.F.; Writing - Original Draft: J.M.V.N and J.G.; Review and Editing: J.M.V.N.; Visualization: J.M.V.N, J.G., I.K.R., C.R.F., Z.G.; Supervision: J.M.V.N. and J.F.; Funding Acquisition: J.M.V.N., J.H., J.F. All authors reviewed and approved of the contents of this manuscript.

---

**Figure S1.**
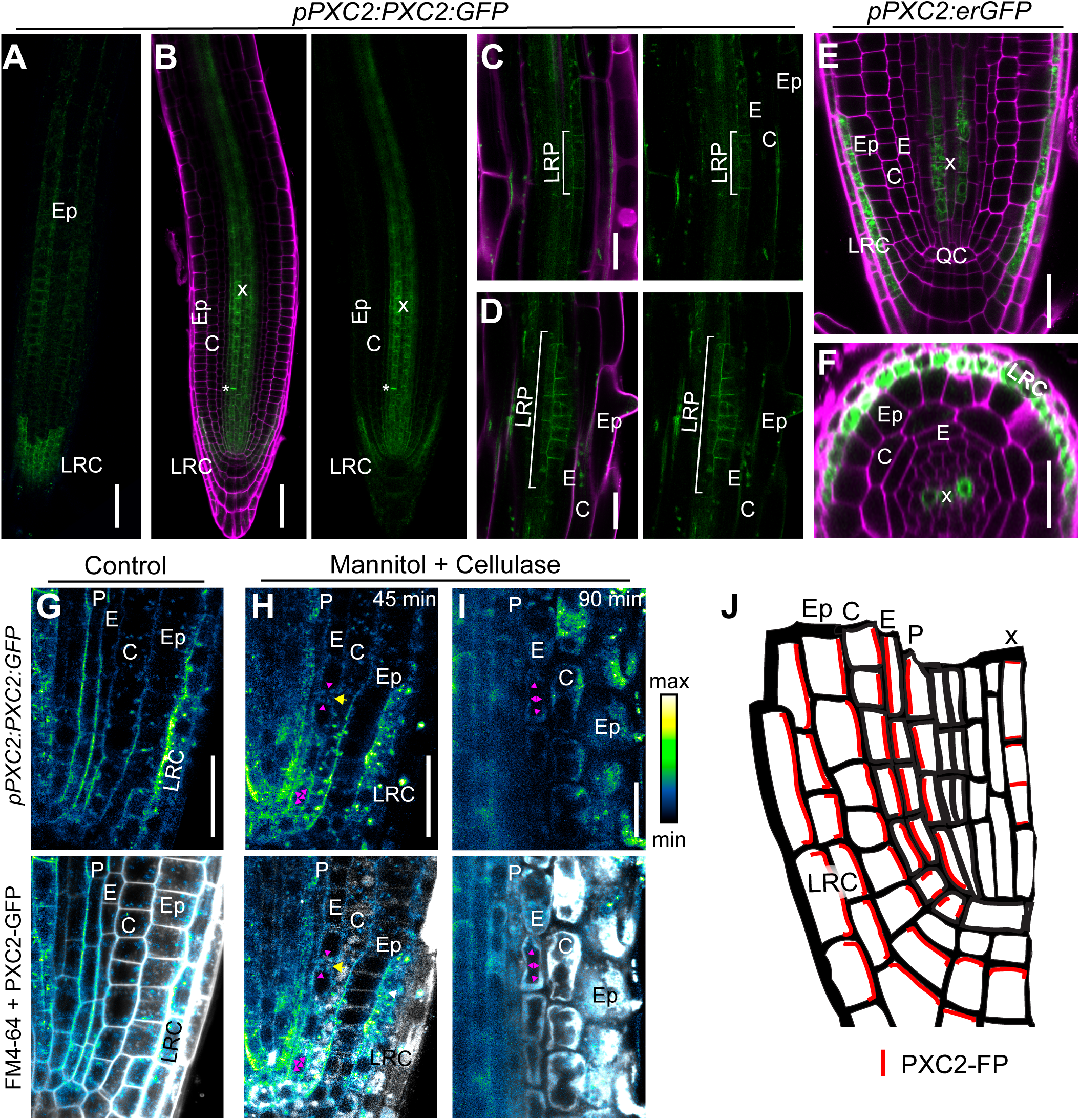
PXC2 promoter activity and PXC2-GFP accumulation in the primary root and lateral root primordia. (A-F) Confocal micrographs of roots stained with propidium iodide (PI, magenta) expressing a GFP reporter (green). (C-F) PXC2-GFP accumulation when driven by pPXC2 (GFP+PI merged and/or GFP alone). (C) PXC2-GFP accumulation in the LRC and epidermis and (D) median longitudinal section of the root tip showing accumulation in xylem cell types, LRC, and new cell plate (asterisk). (E, F) Longitudinal optical sections in the maturation zone showing PXC2-GFP accumulation in early stage lateral root primordia (LRP, brackets). (E) Median longitudinal and (F) transverse optical sections showing pPXC2:erGFP activity in the root tip (GFP+PI merged). (G) Control and (H, I) plasmolysis with partial cell wall degradation in roots expressing pPXC2:PXC2:GFP with GFP alone (upper, intensity color scale) and GFP + FM4-64 (white, merged in lower panels) showing PXC2-GFP accumulation in most cell types, including the endodermis (magenC,DE, Fta arrowheads). Yellow arrow indicates a cell wall between adjacent endodermal cells. (J) Schematic interpretation of PXC2-FP plasma membrane accumulation in root cell types based on endogenous and misexpression (see Fig. 1). Scale bars: (A-F) 25μm, (G-I) 20µm. Abbreviations: (QC) quiescent center, (E) Endodermis, (C) Cortex, (Ep) Epidermis, (LRC) Lateral Root Cap, (P) pericycle, (x) xylem axis.

**Figure S2.**
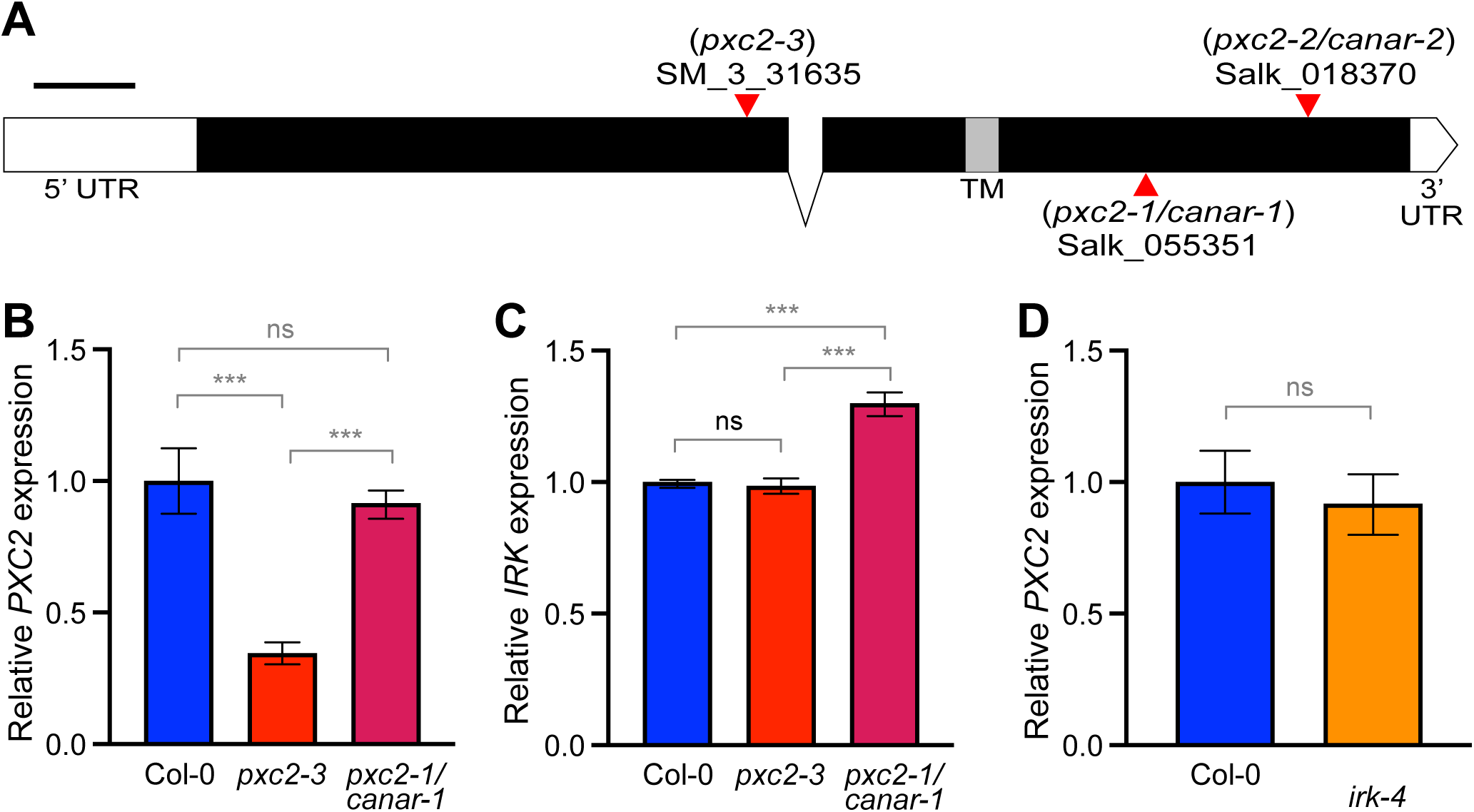
PXC2 gene structure and expression analyses. (A) PXC2 (At5g01890) gene structure with exons as boxes with the coding region in black and the intron as a line. Note that the region encoding the transmembrane (TM) domain is highlighted in gray. The positions of insertional alleles indicated by red triangles, including *pxc2-3* and two previously characterized alleles *pxc2-1/canar-1* and *pxc2-2/canar-2*. Scale bar: 250 base pairs. (B-D) Expression of *PXC2* and *IRK* relative to controls. Data shown from a single biological replicate (of three with similar results) with error bars indicating standard deviation among three technical replicates. Primers for qRT-PCR on *PXC2* are located on either side of the intron downstream of *pxc2-3* insertion site (Table S1). Abbreviations: UTR = untranslated region, ns = not statistically significant, *** = p-value <0.001, Student’s t-test.

**Figure S3.**
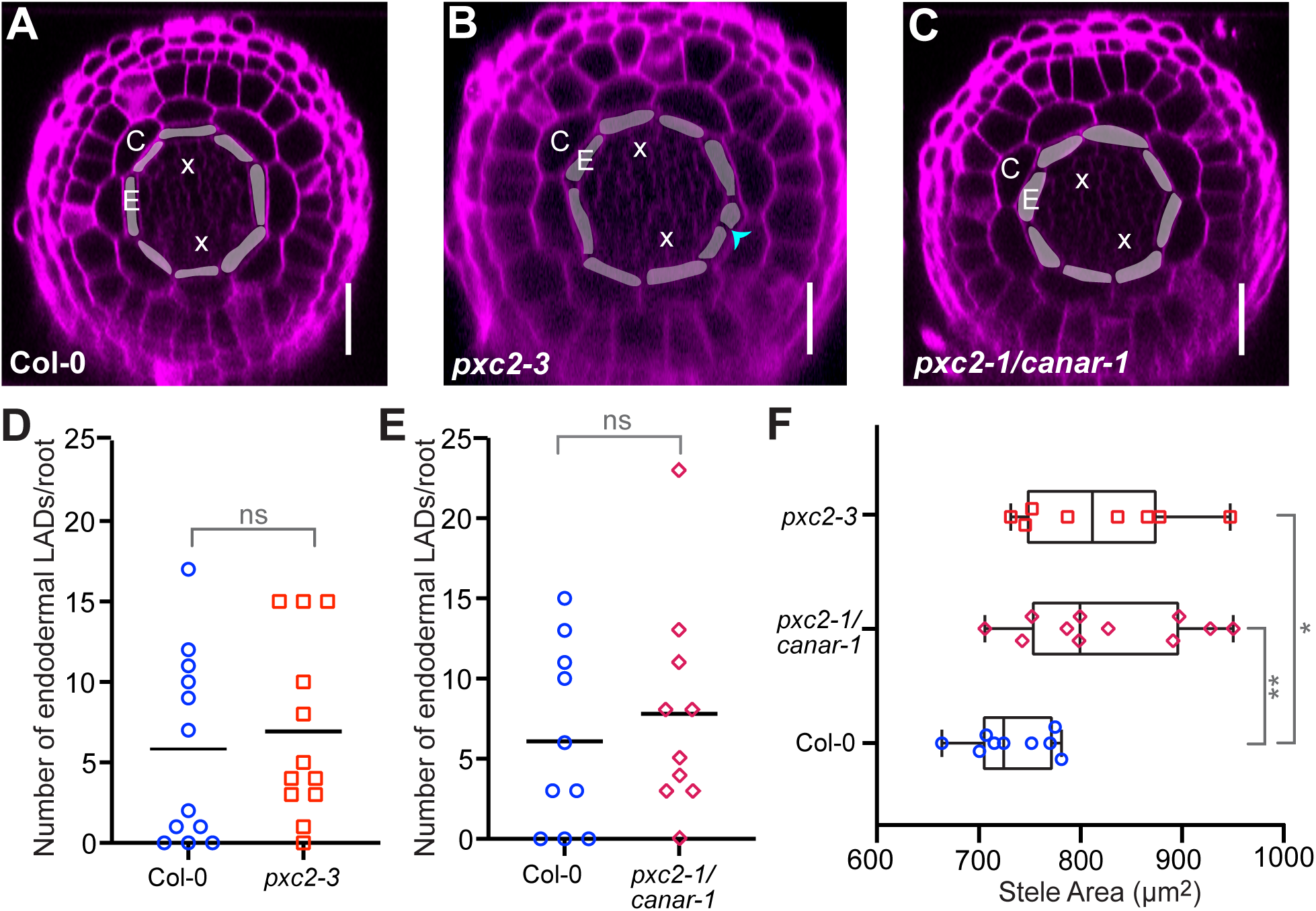
*pxc2-3* and *pxc2-1/canar-1* roots have similar abnormal root phenotypes. (A-C) Confocal micrographs of transverse optical sections from root meristems grown on 0.2x MS, imaged at 6 dps. Roots were stained with PI and endodermal cells are shaded. Cyan arrowhead shows an endodermal LAD. (D-F) Graphs showing (D, E) total number of endodermal LADs per root and (F) stele area in roots at 6 dps, n = 8-12 roots per genotype. Data shown from a single representative biological replicate of ≥3. Note the Col-0 section is repeated from Figure 2, as is the data in (D) with the y-axis rescaled for direct comparison to *pxc2-1/canar-1*. Scatter plots (D, E): Total number of endodermal LADs per root is represented by colored symbols and a black bar indicating the mean for each data set. Box plot (F): whiskers indicate min/max with interquartile range and median shown with black boxes/lines, respectively, with colored symbols indicating measurements of individual roots. Scale bar: 25μm. Abbreviations: C - cortex, E - endodermis, x - xylem axis. Statistics: ns = not significant (p value > 0.5), p values: * = ≤ 0.05, ** = ≤ 0.01, Mann-Whitney test.

**Figure S4.**
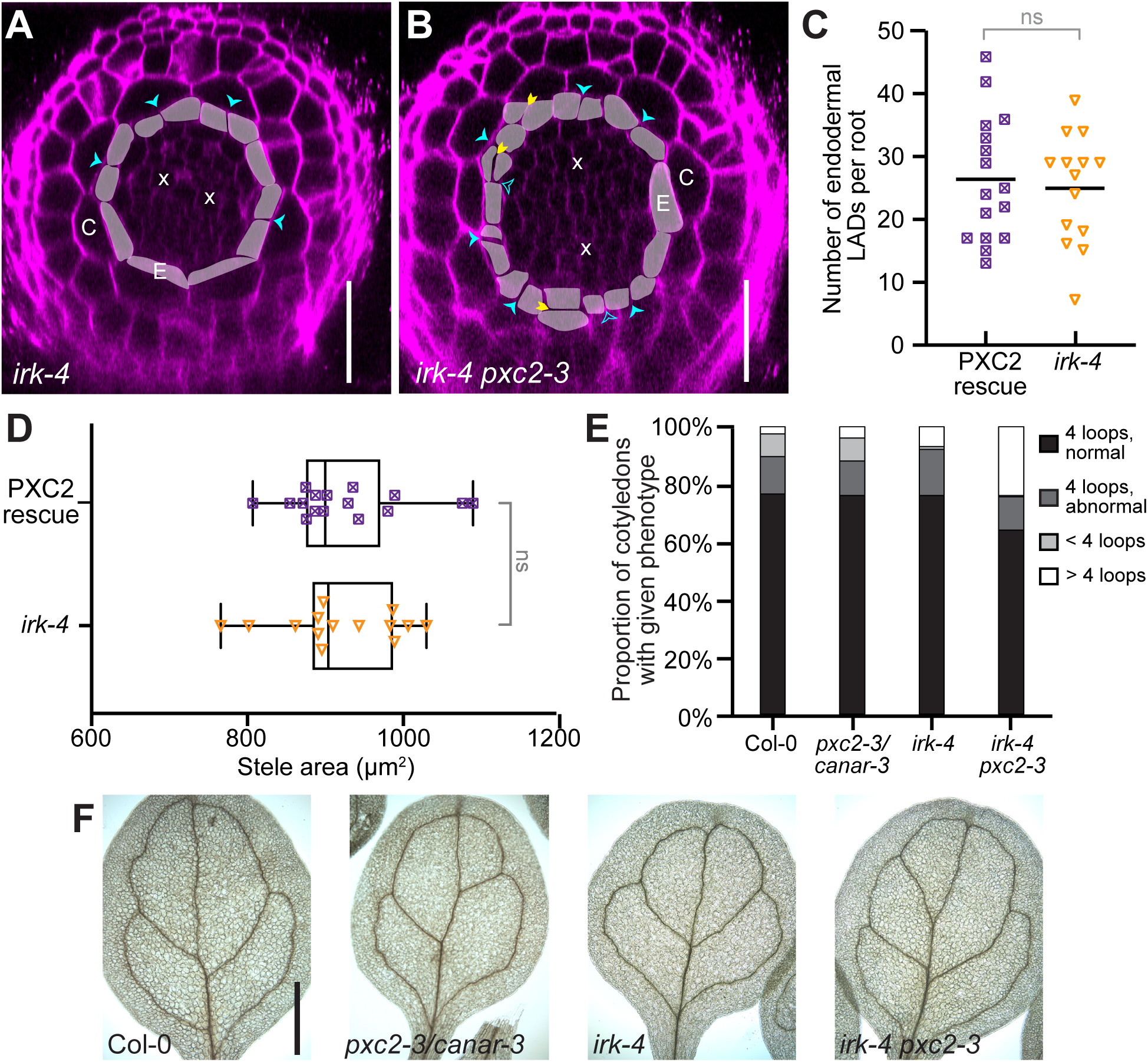
*irk-4 pxc2-3* root and cotyledon vein patterning phenotypes with root phenotype rescued back to *irk-4* by expression of *pPXC2:PXC2:GFP*. (A, B) Confocal micrographs of transverse optical sections from root meristems stained with PI (A) *irk-4* and (B) *irk-4 pxc2-3* at 6 dps with endodermal cells shaded. Cyan arrowheads indicate LADs, yellow arrowheads indicate periclinal divisions, and unfilled cyan arrowheads indicate possible endodermal LADs based on cell position. Scale bars: 25 μm. Note the double mutant phenotype is more severe at 6 dps than 4 dps (compare to Figure 2F). (C, D) At 4 dps, expression of *pPXC2:PXC2:GFP* in *irk-4 pxc2-3* (PXC2 rescue, n = 16) restores the (C) endodermal LAD and (D) stele area phenotypes back to the *irk-4* (n = 14) phenotype. Data from a single representative biological replicate of ≥3. Scatter plot (C): Data for individual roots indicated by colored symbols with the black bar indicating the median. Box plot (D): whiskers indicate min/max with interquartile range and median shown in black boxes/lines, respectively, and colored symbols indicate measurements for individual roots. Error bars = standard deviation. Abbreviations: C - cortex, E - endodermis, x - xylem axis, ns = not statistically significantly different (assayed by Mann-Whitney tests). (E, F) Cotyledon vein pattern phenotypes in various genotypes. (E) Bar graph of quantification of cotyledon vein pattern phenotypes (n = 200 per genotype). (F) Brightfield images of cotyledons showing representative cotyledon vein patterns. Scale bar: 0.5 mm.

**Figure S5.**
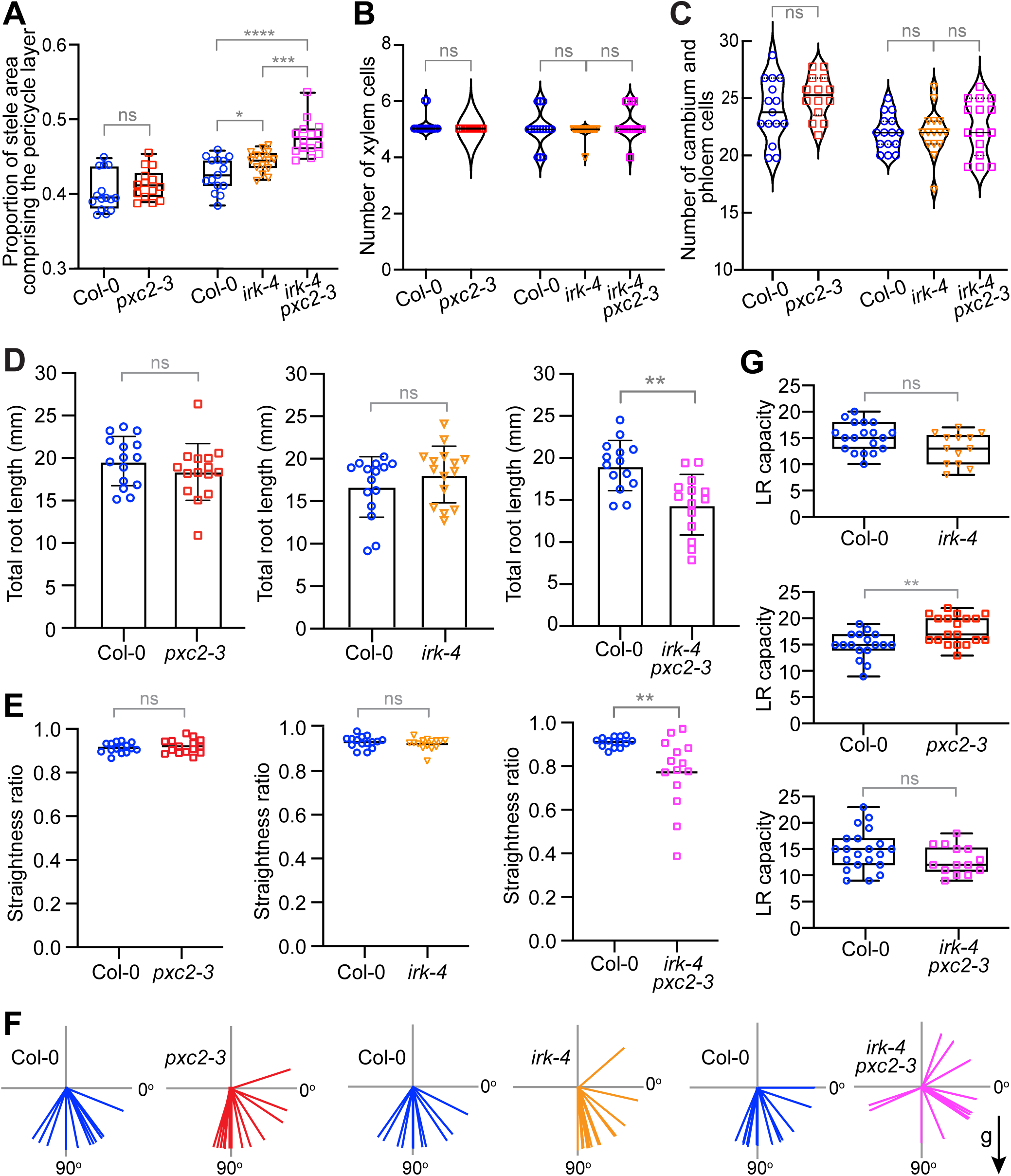
Stele cell and root growth phenotypes in *pxc2-3* and *irk-4* single and double mutants. (A) Box plot showing the proportion of stele area made up of the pericycle layer (n = 14-15 roots per genotype). Whiskers indicate min/max with interquartile range and median shown in black boxes/lines, respectively, and symbols show measurements for individual roots. (B, C) Violin plots showing cell numbers within the stele in one optic section of the root meristem (same roots examined as in A). Symbols show measurements for individual roots with the median and quartiles, shown as solid or dotted black lines, respectively. Data shown are from a single representative biological replicate of ≥ 3 with similar results. (D-G) Graphs showing pairwise comparisons of root growth and phenotypes in Col-0 and *pxc2-3, irk-4*, or *irk-4 pxc2-3* roots at 7 dps (n = 15 per genotype). (D) Total root length where the bar graph shows average length, error bars = standard deviation. (E) Straightness ratio where scatterplot shows data for individual roots as colored symbols and the median indicated by the black bar. (F) Root tip angle 8h after gravistimulation by turning plates 90°. Colored lines = angle of individual root tips. New gravity vector (90°) indicated by g and black arrow, previous gravity vector aligned with 0°. Note the data shown for Col-0 and *irk-4 pxc2-3* in D-F are replicated from Figure 3 for ease of comparison. (G) Box plot showing lateral root (LR) capacity in pairwise comparisons between Col-0 and the mutant genotypes (n = 13-21 per genotype). Whiskers indicate min/max with interquartile range and median shown in black boxes/lines, respectively, and symbols show measurements for individual roots. All data shown is from a single biological replicate of ≥3, with all having similar results. Statistics: ns = not significantly different, p values: * = ≤ 0.05, ** ≤ 0.005, *** = 0.0001, **** = ≤ 0.0001, assayed by Mann-Whitney tests.

**Figure S6.**
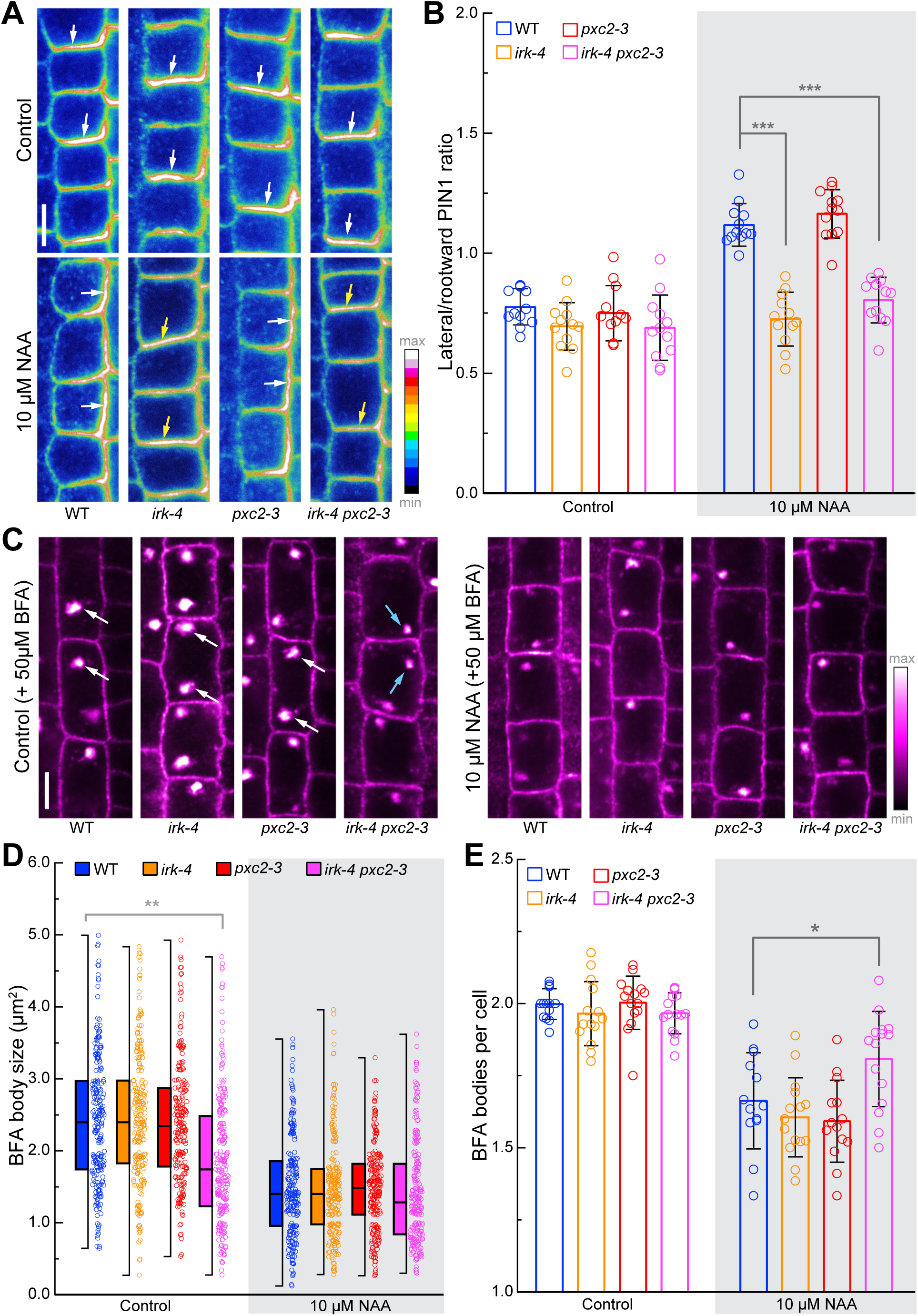
IRK and PXC2 participate in auxin-mediated feedback on PIN1 accumulation and trafficking in roots. (A-B) Auxin-induced PIN1 lateralization of endodermal cells in the longitudinal axis of the root meristem in WT, *irk-4*, *pxc2-3*, and *irk-4 pxc2-3*. (A) *in situ* immunolocalization showing PIN1 in endodermal cells shown with intensity color scale. White arrows show primarily rootward accumulation of PIN1 in all genotypes in control conditions (upper panels) and a shift to lateral accumulation of PIN1 in WT and *pxc2-3* upon auxin treatment, while accumulation remains rootward in *irk-4* and *irk-4 pxc2-3* (yellow arrows). Scale bar = 5 μm. (B) Quantification of PIN1 accumulation as a ratio of signal intensity at the inner lateral versus rootward plasma membrane domains. Bar height indicates the mean ± standard deviation (SD) with individual data points showing the average ratio of different cells in one root (n = >12 roots and at least 200 cells per genotype. Statistics: p-values indicated by asterisks, *** p ≤ 0.001, assayed by a Tukey’s multiple comparison test. (C-E) Auxin-induced PIN1 trafficking with BFA treatment in pericycle cells of WT, *irk-4*, *pxc2-3* and *irk-4 pxc2-3*. (C) PIN1 on the cell membrane (magenta) and accumulation in BFA bodies (white). In control conditions, BFA-bodies of typical size (white arrowheads) and smaller BFA-bodies (blue arrowheads) in *irk-4 pxc2-3* in control conditions. Scale bar = 5 μm. (D-E) Quantification of BFA-body size and BFA bodies per cell (density, n = >220 cells from at least 14 roots per genotype per condition). (D) Half box plots with interquartile range and median shown as black boxes/lines and whiskers showing max/min values. Open circles represent the sizes of individual BFA bodies. (E) Bar graph shows the mean ± SD with open circles indicating measurements for individual roots. Statistics: p-values indicated by asterisks, * p ≤ 0.05 and ** p ≤ 0.01 as determined by Tukey’s multiple comparison test.

**Figure S7.**
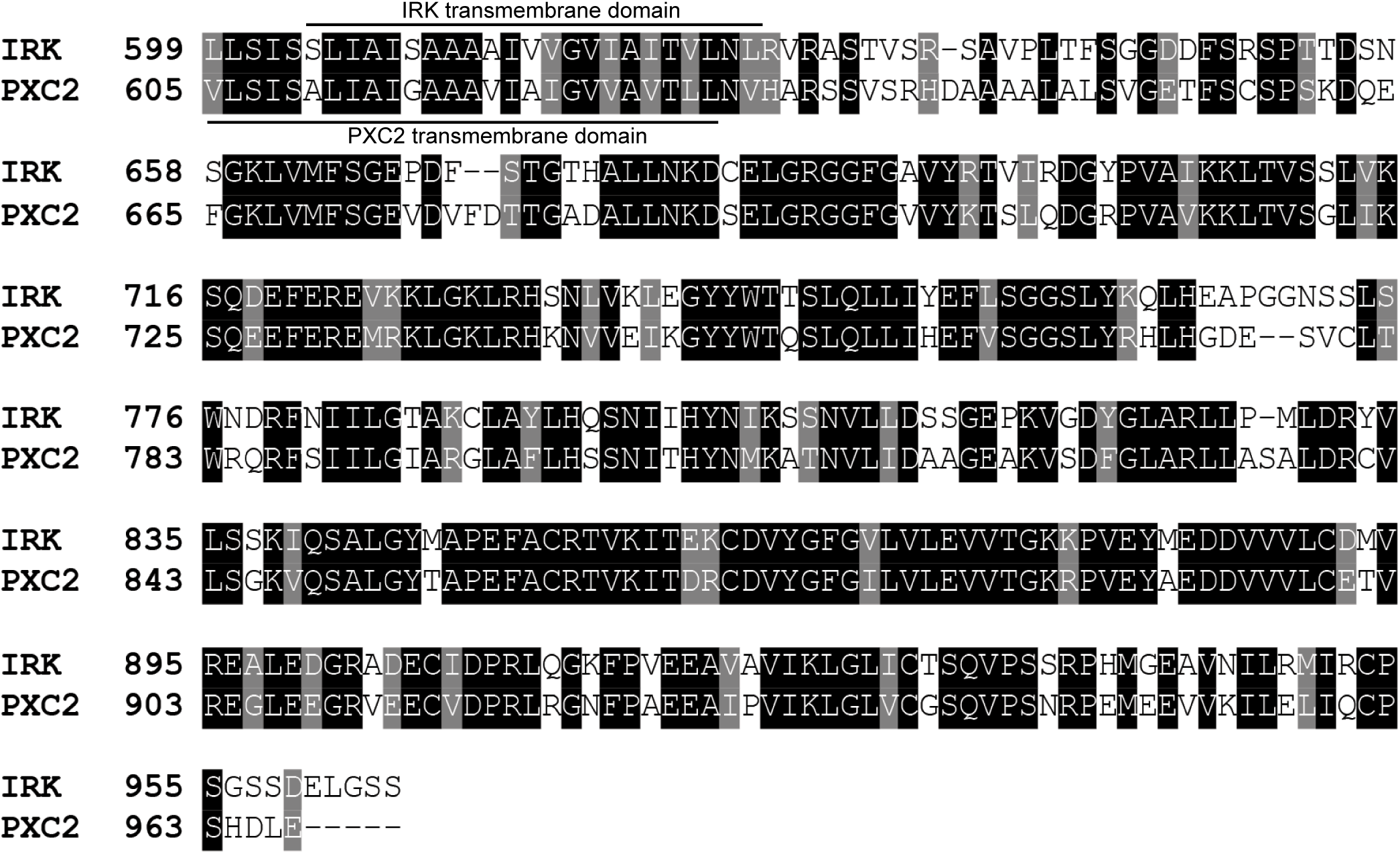
Alignment of IRK and PXC2 intracellular domains. Amino acid alignment begins at the transmembrane domain (marked with the bar). Black shading indicates identical residues and gray indicates similar residues with gaps indicated by dashes.

**Table S1:**
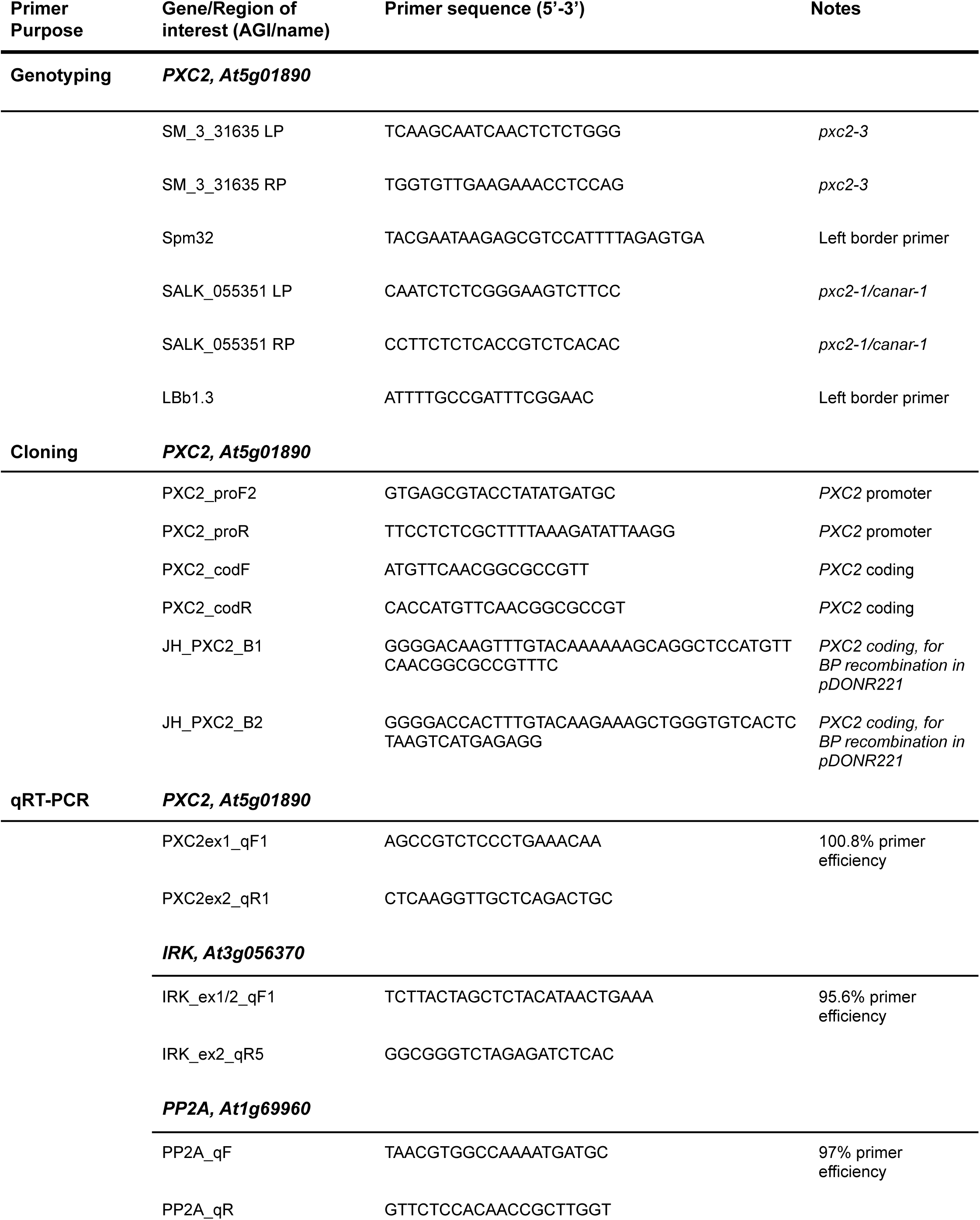
List of primers used in this study.

